# Verbal versus Nonverbal Processing Leads to Generalized Hemispheric Laterality Effects that Span Multiple Networks

**DOI:** 10.64898/2025.12.15.694432

**Authors:** Wendy Sun, Lauren M. DiNicola, Randy L. Buckner

**Affiliations:** Div. of Medical Sciences, Harvard Medical School, Boston, MA; Dept. of Psychology, Center for Brain Science, Harvard University, Cambridge, MA; Dept. of Psychology, University of Virginia, Charlottesville, VA; Dept. of Psychiatry, Massachusetts General Hospital, Boston, MA; Athinoula A. Martinos Center for Biomedical Imaging, Massachusetts General Hospital, Charlestown, MA

## Abstract

Precision neuroimaging was used to explore specialization for verbal versus nonverbal processing. Consistent with prior findings, individuals exhibited a spatially left-lateralized association language network, with regions in both hemispheres robustly responding to processing of meaning-based sentences. We next examined differential responses to verbal (words) versus nonverbal (faces) materials within the same working memory task. The right hemisphere components of the language network responded more strongly to nonverbal than to verbal materials, splitting the network’s functional profile between the hemispheres. Similar patterns were observed across multiple association networks including putative cognitive-control, action-mode, and attention networks. The hemispheric laterality effect was prospectively replicated in a second independent study. These findings highlight a generalized laterality phenomenon that transcends the specialization of individual networks that has been the recent focus of the human systems neuroscience field, and aligns with a broad mechanism that modulates processing between the hemispheres.

## Introduction

A central principle of human brain organization is that the two hemispheres are functionally specialized for verbal versus nonverbal processing. More than a century ago, Paul Broca noted that frontal lesions causing aphasia tend to be on the left (Broca, 1865; see Berker, Berker, and Smith, 1986). Comprehensive explorations of patients with unilateral strokes and patients undergoing unilateral sodium amytal injections (the Wada procedure; Wada and Rasmussen, 1960) reinforced this foundational observation (e.g., Branch, Milner, and Rasmussen, 1964; Raja Beharelle et al., 2010; Sinanović et al., 2011). Additional findings further revealed that distinct functions are right-hemisphere dominant including certain forms of nonverbal processing and spatial attention (e.g., Milner, 1971; Sackeim et al., 1982; Mesulam, 1999; Zatorre, Belin, and Penhune, 2002). Patients with “split-brain”, whose hemispheres are disconnected due to surgical transection of the corpus callosum, reveal that distinct information processing functions can be supported, to a degree, by each hemisphere independent of the other (Gazzaniga, Bogen, and Sperry, 1962; Sperry, 1974; Gazzaniga, 2000).

Building from these seminal discoveries, the present work explored the basis of hemispheric specialization from a systems neuroscience framework. Higher-order cognitive functions are supported by distributed association networks that are made up of anatomically connected brain regions (Geschwind, 1965; Goldman-Rakic, 1988; Mesulam, 1990; 1998). In the human, several of these networks are prominently lateralized, meaning their component regions include greater representation in one hemisphere more than the other. For example, meaning-based language functions are supported by a specialized left-lateralized association network that includes regions at or near classically-defined Broca’s and Wernicke’s areas, as well as distributed regions in superior frontal and inferior temporal cortex (Fedorenko et al., 2010; Fedorenko, Duncan, and Kanwisher, 2012; Braga et al., 2020; Du et al., 2024; see Fedorenko, Ivanova, and Regev, 2024 for review). This association network responds preferentially to processing meaningful sentences, distinguishing it from nearby regions belonging to adjacent association networks (Fedorenko, Duncan, and Kanwisher, 2012; 2013; Braga et al., 2020; Du et al., 2024).

Critically, even though the association language network is strongly left lateralized, there are smaller component regions in the right hemisphere that are part of the network (e.g., see Braga et al., 2020; see also Lipkin et al., 2022). These right-lateralized regions, like their larger left hemisphere counterparts, are robustly activated during meaning-based language processing, indicating that the entirety of the distributed network responds together (see Fig. 21 in Braga et al., 2020, Fig. 1 in Lipkin et al., 2022, extended data Fig. 8 in Malik-Moraleda et al., 2022, and Supplementary Figs. 13 & 14 in DiNicola, Sun, and Buckner, 2023). That is, while the language network regions on the right are smaller than those on the left, they are also reliably activated during meaning-based language processing.

**Figure 1.**
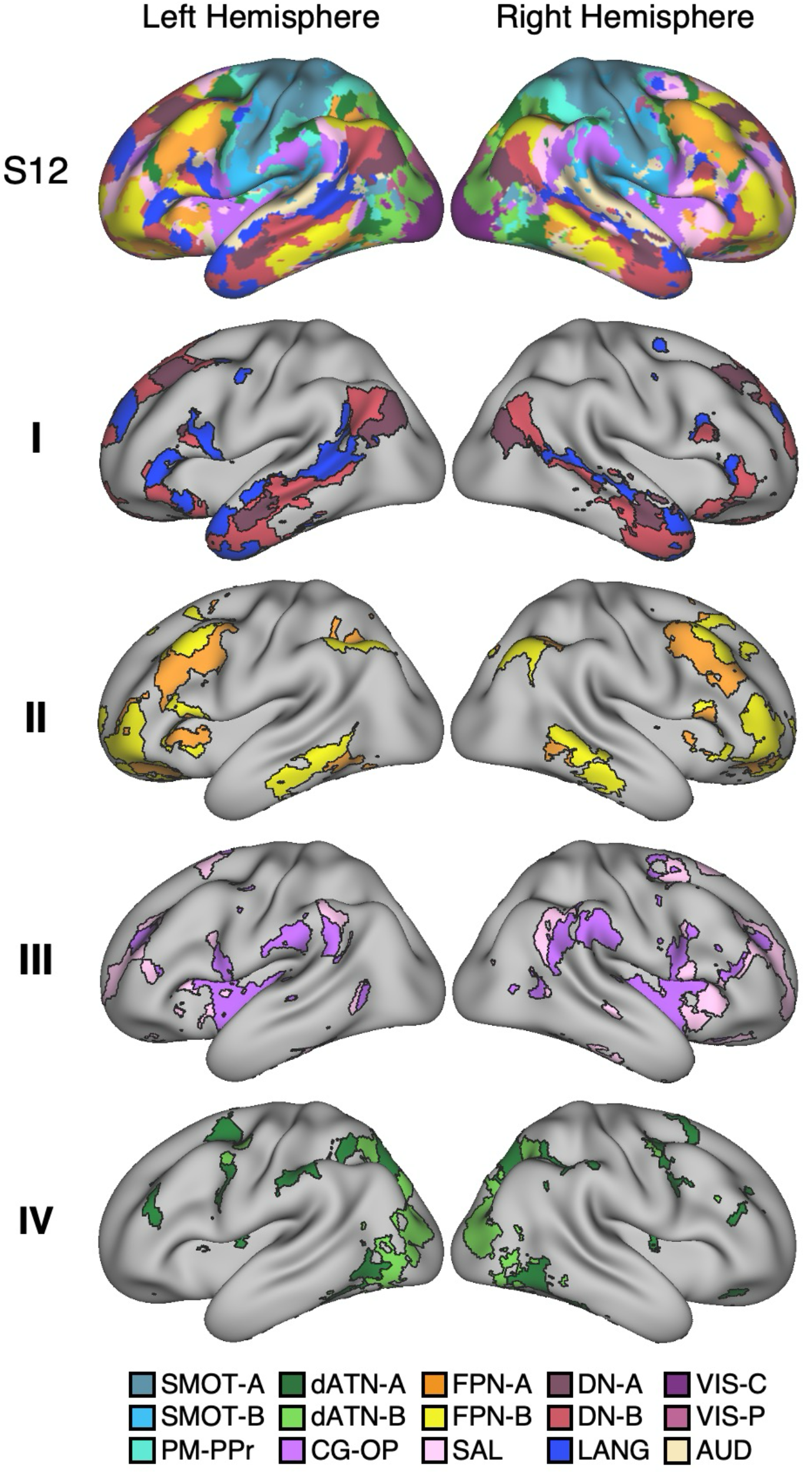
All association networks are bilateral with varied degrees of asymmetry. Network estimates are illustrated for a representative individual from the discovery study (*S12*). The top row shows the anatomical topography of all 15 networks; rows **I**-**IV** highlight the four network groups examined. Note that all networks are bilateral, including the strongly lateralized networks LANG (left lateralized) and FPN-B (right lateralized). **Figure S2** shows the quantitative lateralization for each network from the discovery sample of 14 individuals. DN-A, Default Network A; DN-B, Default Network B; LANG, Language; FPN-A, Frontoparietal Network A; FPN-B, Frontoparietal Network B; CG-OP, Cingulo-Opercular; SAL, Salience; dATN-A, Dorsal Attention A; dATN-B, Dorsal Attention B.

Extending from the finding that there exists a widely distributed left-lateralized association language network, we explored the possibility that functional specialization might arise from distinct networks that differentially occupy the right hemisphere. We initially hypothesized that a specific right-lateralized association network, referred to as Frontoparietal Network B (FPN-B), would be active during processing of nonverbal materials^1^. To anticipate our results, this is not what we found. Rather, across two studies analyzing independent data from intensively sampled individuals, we found that processing of verbal versus nonverbal materials modulates activity between the left and right hemispheres, transcending individual specialized networks and splitting their response properties between the hemispheres. Such results are consistent with a broad, modulatory mechanism that operates between the hemispheres above and beyond the specialization of networks that the field has charted to date.

## Results

### All Association Networks Are Bilateral With Varied Degrees of Asymmetry

Networks were estimated for every individual using a 15-network multisession hierarchical Bayesian model (MS-HBM; Du et al., 2024), and model-free seed-region based correlations confirmed the network estimates. The present analyses focus on nine association networks, organized into four groups based on anatomical features and functional properties (labeled **I**-**IV** in **Figure 1**). Group **I** networks consist of Default Network A (DN-A), Default Network B (DN-B), and Language (LANG). DN-A, DN-B, and LANG are each widely distributed networks that are juxtaposed with one another across the cortical association zones of each hemisphere. Group **II** consists of Frontoparietal Network A (FPN-A) and FPN-B. Both FPN-A and FPN-B include regions that are themselves juxtaposed but distinct from the DN-A / DN-B / LANG cluster (Du et al., 2024). Group **III** consists of the Cingulo-Opercular (CG-OP) network (also termed the “Action-Mode Network”; see Dosenbach, Raichle, and Gordon, 2025 for review), and the spatially similar Salience (SAL) network (see Seeley, 2019 for review). CG-OP expands outwards from motor and somatosensory cortex and includes the anterior insula. SAL abuts CG-OP in several locations but does not surround motor and somatosensory regions. Finally, group **IV** consists of Dorsal Attention Network A (dATN-A) and Dorsal Attention Network B (dATN-B), which juxtapose retinotopic visual regions and include frontal regions at or near to the frontal eye field (FEF; see Corbetta and Shulman, 2002 for review).

All estimated networks included regions in both hemispheres but with varied degrees of lateralization. To quantify the relative proportion of each network’s representation in each hemisphere, the percentage (%) of vertices in each network was calculated for both left and right hemispheres. **Figure S2** shows the mean results across discovery dataset participants. As expected, LANG was left lateralized (higher % of vertices on the left; Braga et al., 2020; Lipkin et al., 2022) and FPN-B was right lateralized (higher % of vertices on the right; Braga et al., 2020). Critical to our current explorations, all networks had components in both hemispheres (**Figure S4**).

### Bilateral Components of Each Association Network Respond When Activated in a Task-Specific Manner

We next characterized the functional engagement of the networks in task data within the same individuals. These task data were completely independent from the data used to estimate networks. **Figure 2** shows the hemisphere-specific responses in each network for the Sentence Processing and N-Back Load Effect task contrasts. Consistent with prior studies, LANG was recruited by the Sentence Processing task contrast. Despite LANG being left lateralized, both hemispheres of the network were robustly and preferentially activated by the Sentence Processing task contrast. In the N-Back Load Effect task contrast, both hemispheres of FPN-A and dATN-A were recruited, and to a lesser degree FPN-B and SAL. These results confirm, as expected, that these two distinct task contrasts differentially activate specific networks. The responses are bilateral, including both the left and right hemisphere components of the networks.

**Figure 2.**
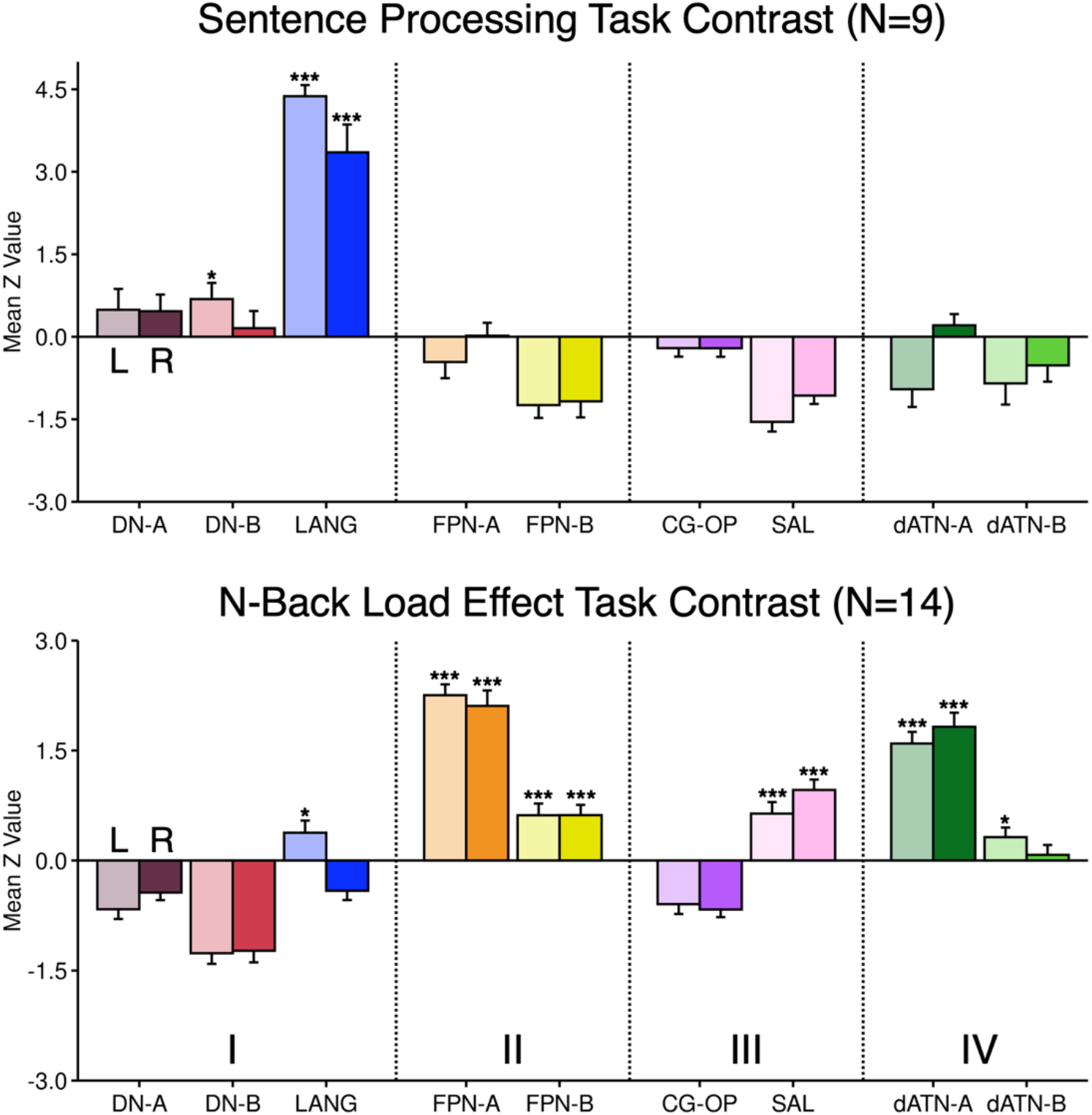
Association networks respond bilaterally in a task-specific manner. The mean response is displayed for all participants from the discovery study for each of the association networks, separately for the left (L) and right (R) hemisphere regions. Lighter-shaded bars represent the left hemisphere, and darker-shaded bars represent the right hemisphere. Error bars show standard error of the mean. (**Top**) Both hemispheres of the LANG network are robustly activated by the Sentence Processing task contrast, even though the network regions in the left hemisphere are larger than those on the right (see **Figure S2**). (**Bottom**) Both hemispheres of FPN-A and dATN-A are robustly activated by the N-Back Load Effect task contrast, with more modest responses in FPN-B and SAL. Asterisks indicate significant positive responses. * = *p* < 0.05, ** = *p* < 0.01, *** = *p* < 0.001.

### Multiple Association Networks Show a Lateralized Response When Processing Nonverbal Materials

The effect of verbal versus nonverbal processing was examined within the N-Back task by directly contrasting the blocks using face stimuli to those using word stimuli (the N-Back Face > Word task contrast). What emerged was surprising: many networks, across all network groupings, showed differential responses between the hemispheres for the N-Back Face > Word task contrast (**Figure 3**).

**Figure 3.**
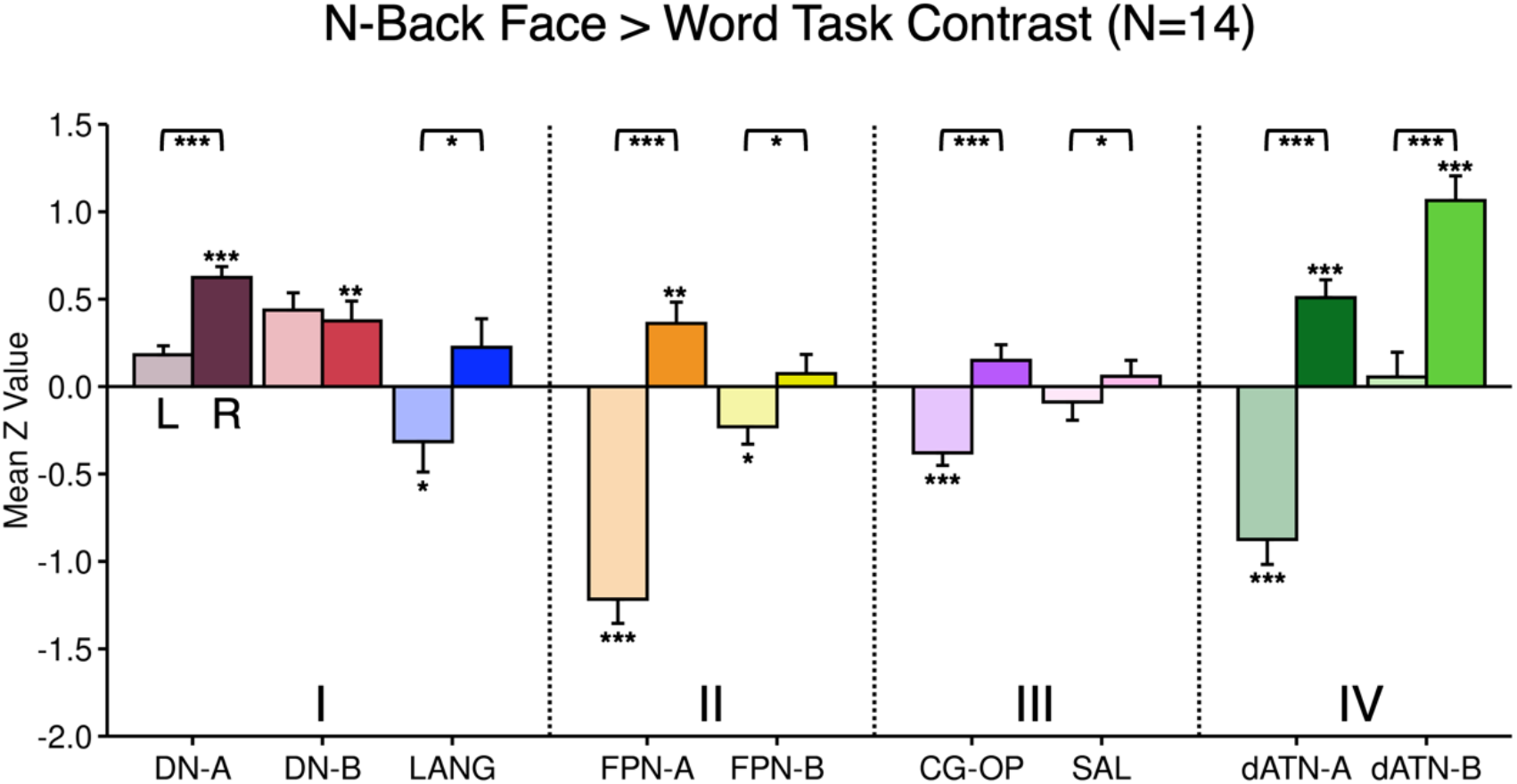
Association networks show lateralized responses for verbal versus nonverbal processing. The mean response for the N-Back Face (nonverbal) > Word (verbal) task contrast is displayed for each of the association networks. Lighter-shaded bars represent the left (L) hemisphere, and darker-shaded bars represent the right (R) hemisphere. Error bars show standard error of the mean. Laterality effects are observed across 8 of the 9 networks, such that the left hemisphere of each network is preferentially recruited by verbal materials, while the right hemisphere is preferentially recruited by nonverbal materials. The effect spans all network groups. Bracketed asterisks indicate that the direct comparisons between the left and right hemisphere responses are significantly different. Asterisks by the bars indicate the individual bar’s response is significantly different from zero (positive in the right hemisphere, negative in the left hemisphere). * = *p* < 0.05, ** = *p* < 0.01, *** = *p* < 0.001.

We had originally envisioned that the effect of verbal versus nonverbal processing might elicit differential responses between networks, perhaps with FPN-B responding more to faces as contrast to words (FPN-B is right lateralized and has been challenging to link to a specific function; see Du et al., 2024)^1^. That is not what was observed. The results revealed that many networks display hemispheric laterality effects splitting the right and left components of the networks.

Statistical tests supported this result. A repeated measures ANOVA with hemisphere (left, right) as the within-subject factor and mean z-value as the dependent variable showed a significant main effect of hemisphere (*F*(1,13) = 69.2, *p* < 0.001). Post-hoc paired t-tests within each network demonstrated significant differences between hemispheres for 8 of the 9 networks. That is, functional dissociation between the hemispheres was observed generally across networks, departing from the differential and bilateral co-activation patterns observed in the standard task contrasts used to isolate the task effects. Note that the lateralization effect elicited by the N-Back Face > Word task contrast was present for the strongly lateralized and specialized LANG network as well as the CG-OP network that was not differentially activated by the main task contrasts (see **Figure 2**). This pattern of results was unexpected.

To further explore the N-Back Face > Word task contrast, the maps of the effect were plotted for each individual participant on the cortical surface as well as for the group average (**Figure 4**). While the idiosyncratic spatial details differed between participants, a strong lateralized pattern was evident in every participant: the left hemisphere exhibited broad responsiveness to words, while the right hemisphere demonstrated broad responsiveness to faces. The pattern encompassed extensive regions of association cortex, including regions in frontal, parietal, and temporal cortices. Thus, the N-Back Face > Word task contrast maps also support the observation of a broad laterality effect that spans multiple association networks.

**Figure 4.**
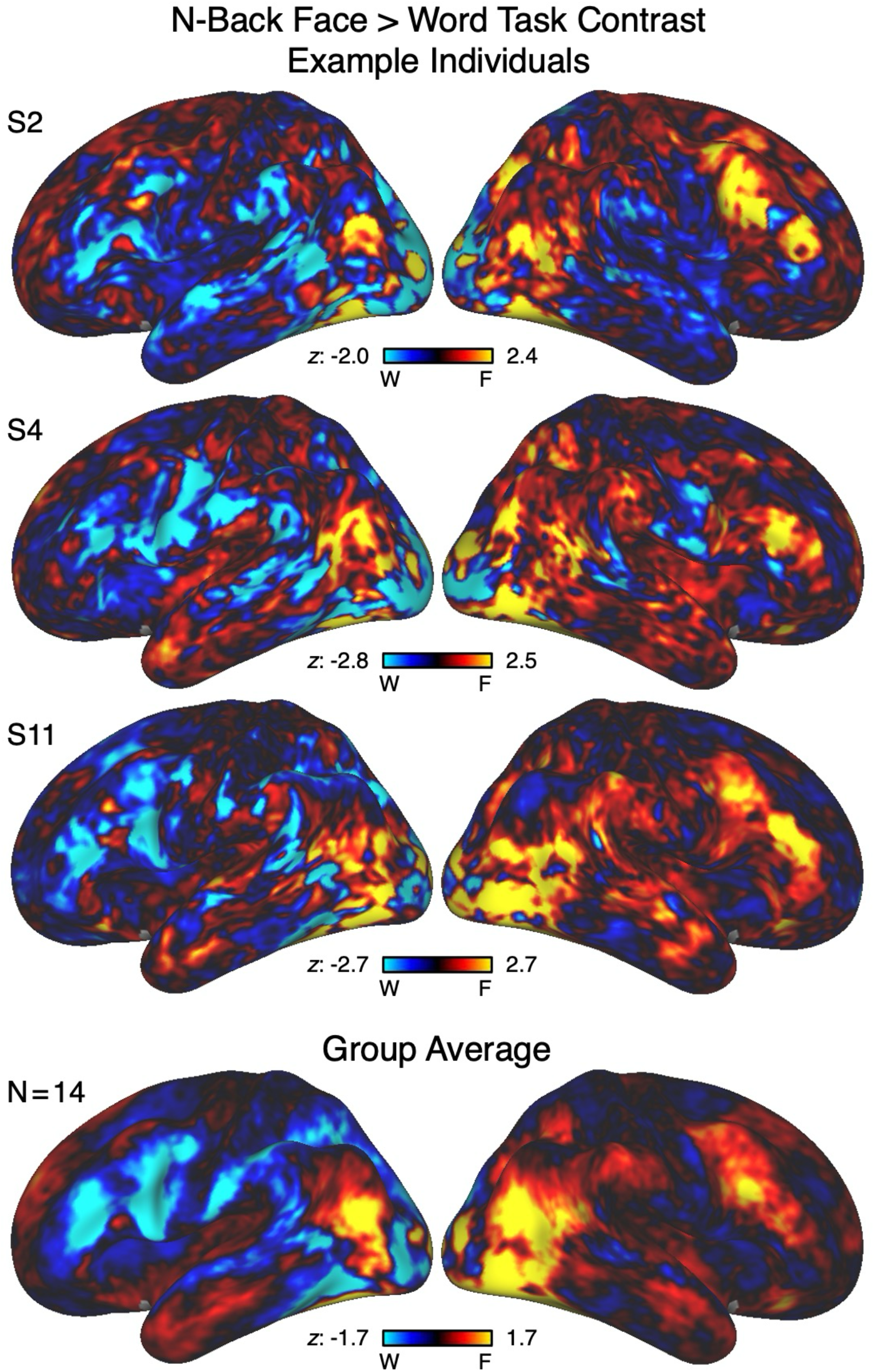
Verbal versus nonverbal laterality effects extend widely across association cortex. Maps of the laterality effect are shown for three representative individuals (**Top Three Rows**) as well as for the average group of participants (**Bottom Row**) from the discovery study. The N-Back Face (nonverbal; red/yellow) > Word (verbal; blue) task contrast is visualized as z-statistical maps on the inflated cortical surface. The data are shown unthresholded to facilitate complete visualization of the effect in each direction. Note that the left hemisphere is broadly responsive to words, and that the right hemisphere is broadly responsive to faces. This distributed pattern spans frontal, parietal, and temporal cortices. W, words; F, faces.

### Generalized Laterality Effects Replicate in Independent Data

To follow up on the surprising results of the discovery study, we turned to replication in the independent data sample. **Figure S1** shows network estimates in one representative participant from the replication study. Group means of the % of vertices in each hemisphere belonging to each network confirmed again that LANG is left lateralized, and FPN-B is right lateralized with all networks having components in both the left and right hemispheres (**Figure S3**). This was true for every participant (**Figure S5**).

As **Figure 5** further reveals, the general task response patterns were similar to the patterns observed in the discovery study (compare **Figure 2** to **Figure 5**). Specifically, component regions of both hemispheres of the LANG network were preferentially recruited by the Sentence Processing task contrast (**Figure 5**), while the N-Back Load Effect task contrast preferentially recruited FPN-A and dATN-A in both hemispheres, as well as FPN-B and SAL to a lesser degree. Once again, CG-OP revealed no response in either of the main task contrasts.

**Figure 5.**
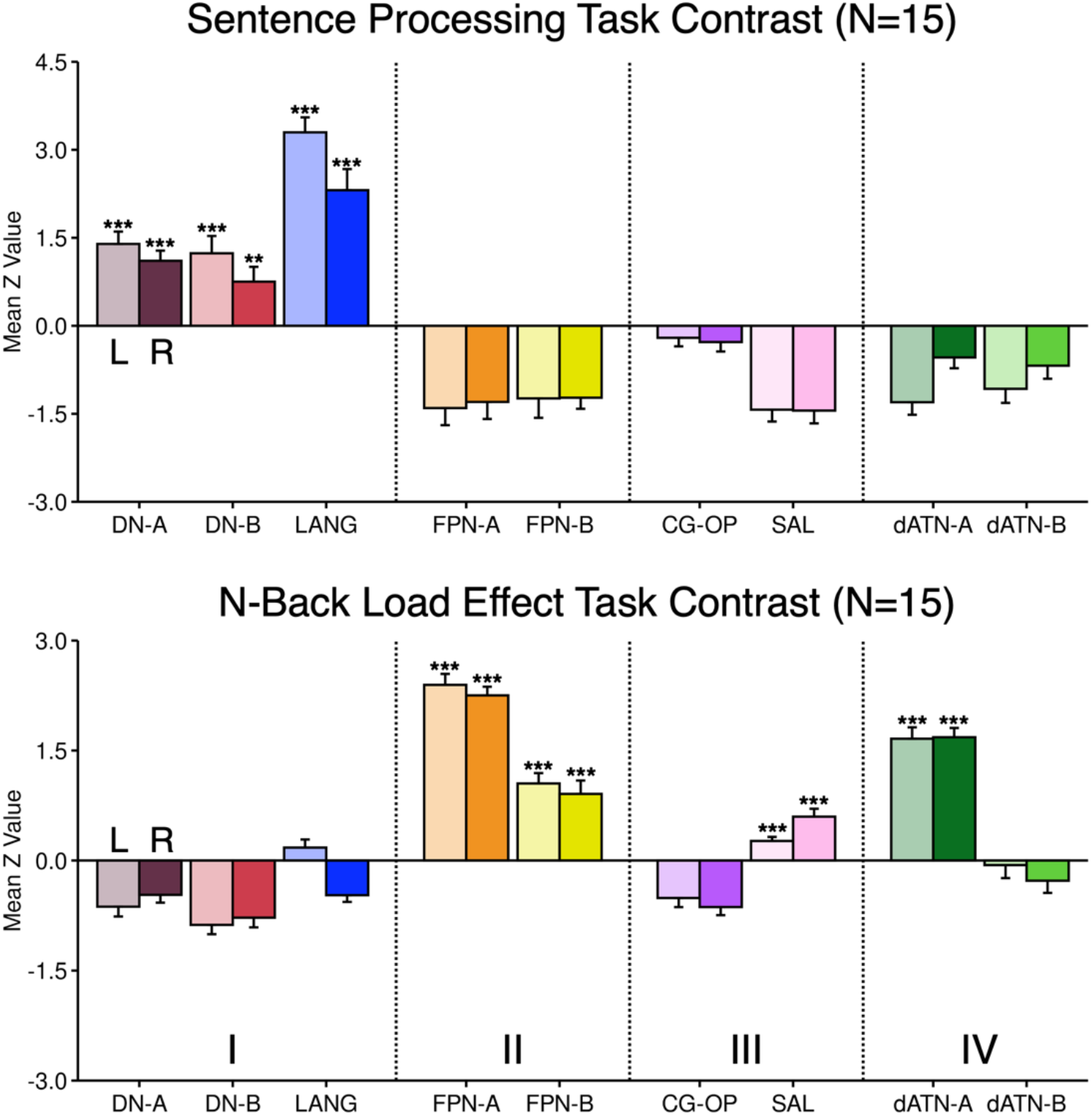
Association networks again respond bilaterally in a task-specific manner. The mean response is displayed for all participants from the replication study for each of the association networks, separately for the left (L) and right (R) hemisphere regions. Lighter-shaded bars represent the left hemisphere, and darker-shaded bars represent the right hemisphere. Error bars show standard error of the mean. (**Top**) Replicating the discovery study, both hemispheres of the LANG network are again activated by the Sentence Processing task contrast, even though the network regions in the left hemisphere are larger than those on the right (see **Figure S3**). (**Bottom**) Both hemispheres of FPN-A and dATN-A are again robustly activated by the N-Back Load Effect task contrast, with more modest responses in FPN-B and SAL. Asterisks indicate significant positive responses. * = *p* < 0.05, ** = *p* < 0.01, *** = *p* < 0.001.

For the critical test, we examined the hemispheric laterality effect in four association networks that span a range of functions chosen *a priori* to test the hypothesis that verbal versus nonverbal processing elicits generalized laterality effects that transcend specific networks. One representative network from each group was chosen: LANG (group **I**), FPN-A (group **II**), CG-OP (group **III**), and dATN-A (group **IV**). Based on the findings from the discovery study, we hypothesized that each network would show a significant N-Back Face > Word task contrast effect. **Figure 6** displays the results.

**Figure 6.**
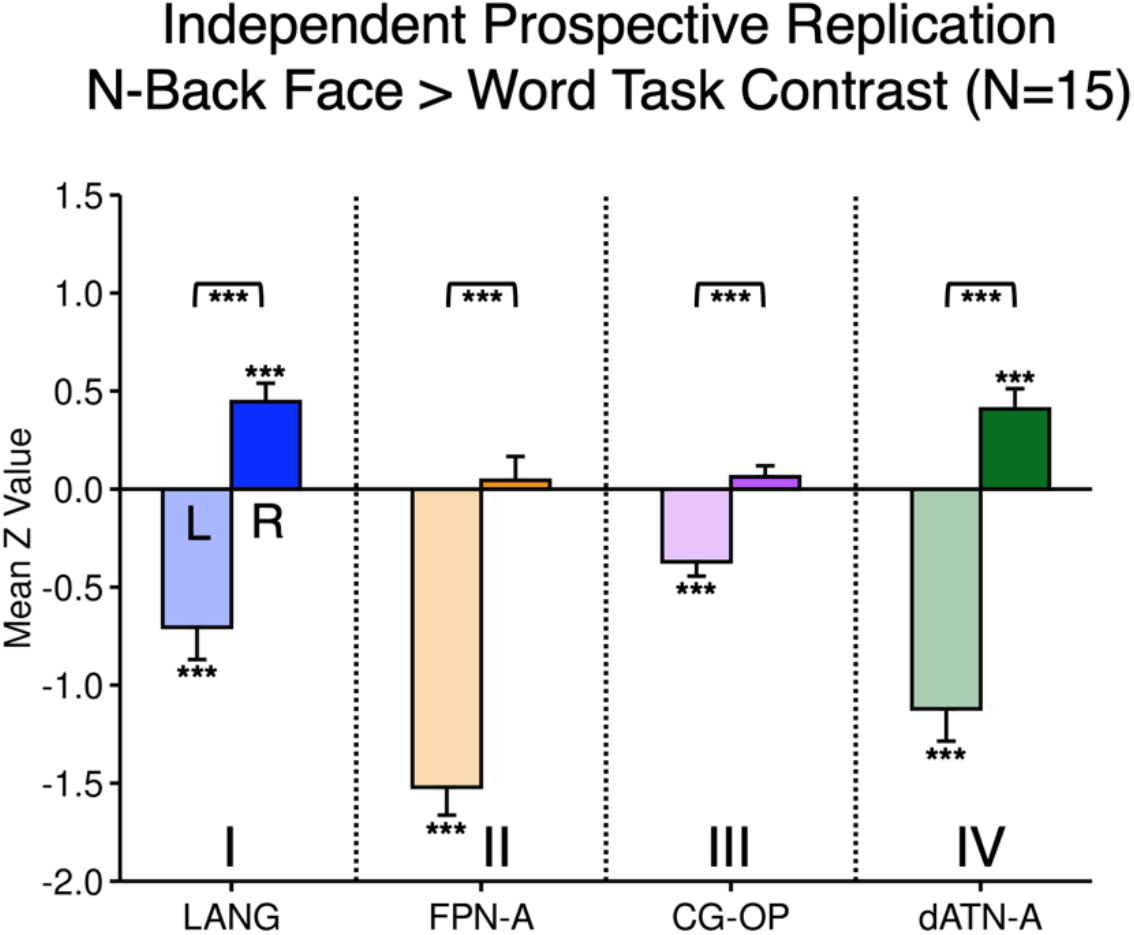
Prospective replication of lateralized responses for verbal versus nonverbal processing. The mean response for the N-Back Face (nonverbal) > Word (verbal) task contrast is displayed for the *a priori* targeted association networks. Lighter-shaded bars represent the left (L) hemisphere, and darker-shaded bars represent the right (R) hemisphere. Error bars show standard error of the mean. The laterality effect was observed in each of the four targeted networks, such that the left hemisphere of each network is preferentially recruited by verbal materials, while the right hemisphere is preferentially recruited by nonverbal materials. The effect spans all network groups. Bracketed asterisks indicate that the direct comparisons between the left and right hemisphere responses are significantly different. Asterisks by the bars indicate the individual bar’s response is significantly different from zero (positive in the right hemisphere, negative in the left hemisphere). * = *p* < 0.05, ** = *p* < 0.01, *** = *p* < 0.001.

The N-Back Face > Word task contrast dissociated the left and right hemispheres across all four tested networks. A repeated measures ANOVA revealed a significant main effect of hemisphere: *F*(1,14) = 219.1, *p* < 0.001. All four planned paired t-tests were significant (**Figure 6**). LANG: *t*(14) = 8.9, *p* < 0.001; FPN-A: *t*(14) = 14.2, *p* < 0.001; CG-OP: *t*(14) = 6.6, *p* < 0.001; dATN-A: *t*(14) = 9.5, *p* < 0.001.

Visualizing the N-Back Face > Word task contrast maps again revealed the hemispheric laterality effect (**Figure 7**), with distributed regions in the left hemisphere recruited for words, and distributed regions in the right hemisphere recruited for faces. The pattern of differential response encompassed broad regions of association cortex, replicating the pattern observed in the discovery study. Thus, all results from both studies – quantitative and qualitative – converge to support the observation of a broad hemispheric laterality effect that transcends individual networks.

**Figure 7.**
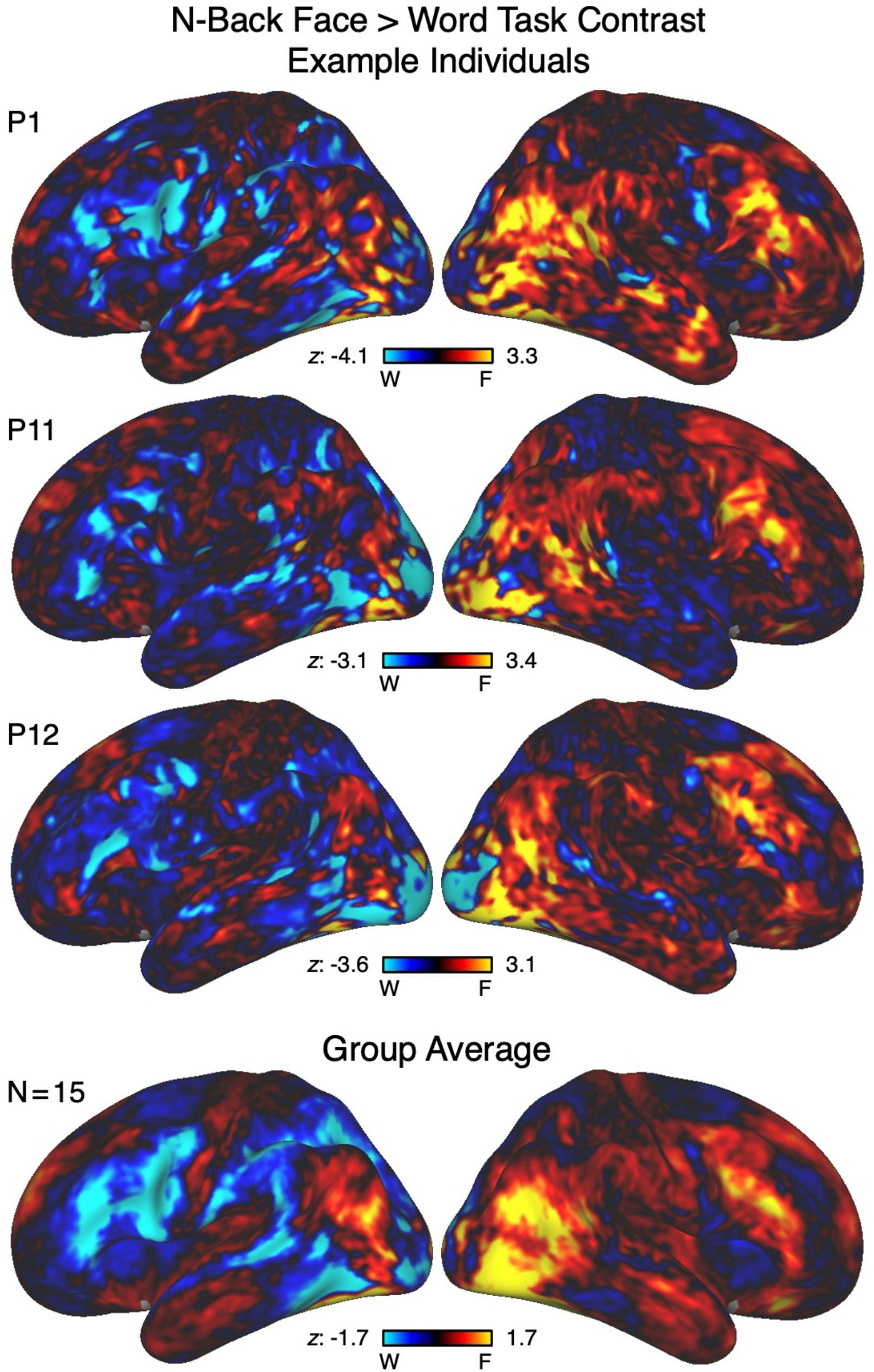
Verbal versus nonverbal laterality effects again extend widely across association cortex. Maps of the laterality effect are shown for three representative individuals (**Top Three Rows**) as well as for the average group of participants (**Bottom Row**) from the replication study. The N-Back Face (nonverbal; red/yellow) > Word (verbal; blue) task contrast is visualized as z-statistical maps on the inflated cortical surface. The data are shown unthresholded to facilitate complete visualization of the effect in each direction. Note that the left hemisphere is again broadly responsive to words, and that the right hemisphere is responsive to faces. W, words; F, faces.

## Discussion

Generalized hemispheric laterality effects were observed across multiple association networks, splitting them such that their right-lateralized components preferentially responded to nonverbal materials. This result is surprising because, in many contexts, association networks activate cohesively, including their bilateral component regions (**Figures 2 and 5**). We found that these same networks can be functionally divided, exhibiting differential hemispheric lateralization during verbal versus nonverbal processing (**Figures 3 and 6**). This pattern suggests the presence of a broad modulatory mechanism capable of driving hemisphere-wide preferential processing that transcends individual specialized networks.

### Nonverbal Processing Leads to Pan-Network Hemispheric Laterality Effects

Association networks have regions in both hemispheres that are recruited together across a variety of task demands. We and others have shown this phenomenon in networks that respond preferentially to domain-specific language (e.g., see Braga et al., 2020; Lipkin et al., 2022) and domain-flexible cognitive control (e.g., see Fedorenko, Behr, and Kanwisher, 2011 and DiNicola, Sun, and Buckner, 2023). Here, we re-demonstrate the bilateral nature of association networks (**Figures 2 and 5**).

Given the bilateral response of association networks, we initially hypothesized that, when presented with stimuli across verbal and nonverbal domains, the networks would respond together with a right-lateralized network (e.g., FPN-B) responding more to nonverbal materials. That is, when considering networks as the functional unit, one might expect that verbal materials would activate both hemispheres of left-lateralized association networks, while nonverbal materials would activate both hemispheres of right-lateralized association networks. The present results indicate this is not the case.

Instead, the lateralized functional regions of multiple networks differentially responded to nonverbal processing. The functional dissociation between the hemispheres was present in spatially discontinuous networks that markedly differed in their functional properties (**Figures 3 and 6**). Prior studies contrasting verbal (word) versus nonverbal (face) processing in group-averaged data have observed hemispheric laterality effects that include robust effects in prefrontal cortex (Kelley et al., 1998; McDermott et al., 1999; Golby et al., 2001; Ballotta et al., 2023). The spatial pattern of right-hemisphere responses in these prior studies was in the vicinity of the FPN-B network, further raising the possibility that differential response in FPN-B might be the origin of the hemispheric laterality effect.

Here we isolated the left and right components of all established association networks within the idiosyncratic anatomy of individual participants and found that contrasting verbal versus nonverbal processing split individual networks between the hemispheres. The pattern could be readily appreciated in the maps of differential Face > Word response in individual participants (**Figures 4 and 7**).

We were especially struck by the hemispheric dissociation of the LANG network in both studies, as this network is the most consistently anatomically left-lateralized association network (e.g., Braga et al., 2020; Fedorenko, Behr, and Kanwisher, 2011; Lipkin et al., 2022). Notably, the hemispheric laterality effect also extended to association networks putatively involved in external attention (dATN-A) and action planning (CG-OP). The laterality effect in CG-OP was present even though this network was not differentially activated in the main task comparisons (**Figures 2 and 5**). Moreover, the response demands were held constant between the verbal and nonverbal conditions. Thus, the hemispheric laterality effect was broad, transcending individual specialized association networks.

### Considerations for Hemispheric Modulation

The discovery of a generalized laterality effect spanning multiple, differentially specialized association networks suggests the existence of a mechanism that broadly modulates activity between the hemispheres. In this light, it is useful to consider other examples of large-scale hemispheric modulation.

A prominent example occurs during unihemispheric slow-wave sleep (USWS). During USWS, one hemisphere rests while the other remains alert for essential ecological functions, such as predator vigilance in birds and surfacing to breathe in cetaceans (Rattenborg et al., 2000). Parietal and occipital electrocorticogram recordings in sleeping cetaceans reveal high-amplitude, low-frequency activity in the sleeping hemisphere and low-amplitude, high-frequency activity in the awake hemisphere. Even in species that do not exhibit USWS, there is evidence for asynchronous hemispheric activity during sleep. For example, Fenk and colleagues (2023) demonstrated that in bearded dragons, rapid eye movement (REM) sleep exhibits a winner-take-all dynamic between the hemispheres. When recording from the left and right claustra during REM sleep, they found that potentials on one side led the other by approximately 20ms, with stronger activity in the leading side and periodic switching between leading and lagging hemispheres. Although it would be premature to directly link these sleep-related phenomena and the present findings in awake humans, it is intriguing to consider that ancient mechanisms may have evolved to modulate activity and arousal levels between the hemispheres.

How might hemispheric modulation arise in a complex system like the human brain? Drawing on dynamical systems theory, Wang and Liu (2020) review a framework of “chimera states” that may support the coexistence of both unified and independent hemispheric processing. Chimera states (Majhi et al., 2019) are proposed to occur when part of a system exhibits coherent (synchronized) activity while another part remains incoherent (desynchronized). One possibility, then, is that dynamic changes in neuronal synchrony and cross-hemispheric communication – via the corpus callosum and commissures – contribute to patterns of neural segregation and hemispheric specialization (see Ocklenburg and Guo, 2024 for a recent review). This framework aligns with our observation of broad hemispheric dissociations during verbal versus nonverbal processing, in which association networks that typically show bilateral co-activation can also functionally modulate by hemisphere under the right conditions.

Relating the theory of chimera states to functional hemispheric specialization may also relate to the seminal research on split-brain patients. Sperry, Gazzaniga and colleagues demonstrated that when cross-hemispheric communication via the corpus callosum is disturbed, each hemisphere can operate with a degree of independence, supporting distinct verbal versus nonverbal processing modes (Gazzaniga, Bogen, and Sperry, 1962; Sperry, 1974; Gazzaniga, 2000). While split-brain studies reveal the brain’s capacity for hemispheric independence under extreme conditions, our findings suggest that similar principles of dynamic hemispheric segregation and integration may operate in more subtle ways in the intact brain.

### Limitations and Future Directions

Despite the robustness of observed functional asymmetries, the precise neurobiological mechanisms governing these hemispheric effects are not directly informed by the present study. Research on “cognitive” chimera states (Bansal et al., 2019) posits that artificially stimulating subcortical regions along the midline might yield a synchronized state across distributed (primary and higher-order) systems in the brain, compared to stimulating other regions. These in-silico experiments imply a global and facilitatory role of subcortical regions in coordinating brain states. Given that association networks have representations in the thalamus (Du et al., 2025; see also Shine et al., 2023 for review), future work could directly explore how subcortical inputs contribute to the generalized laterality effects we observe, for example, in patients with distributed stereo-electroencephalography (sEEG) implants. With opportune cortical and subcortical contacts in both hemispheres, this approach offers high spatiotemporal resolution to directly measure neuronal synchrony and possible chimera state at the mesoscale. Moreover, studying populations with corpus callosum abnormalities, such as patients with callosal agenesis or degeneration, could provide natural experiments to probe the role of cross-hemispheric communication in mediating broad hemispheric dissociations.

Another limitation of the present study is the relatively small set of cognitive tasks. While our verbal and nonverbal N-Back paradigm provides evidence for broad hemispheric dissociations, it remains unclear whether these patterns will generalize to other cognitive processes. Future studies should incorporate a wider range of tasks and stimuli to determine the degree to which our observed laterality effects generalize.

Finally, the possible translational implications of these hemispheric dissociations should be examined. Hemisphere-specific targeting is already important in transcranial magnetic stimulation (TMS) for neuropsychiatric disease. A common clinical target for Major Depressive Disorder (MDD) is left dorsolateral prefrontal cortex (dlPFC; see Perera et al., 2016 for review), with emerging precision approaches enabling the targeting of distinct functional networks defined in the individual (Lynch et al., 2022). In some cases, the right dlPFC is targeted for anxiety and trauma-related disorders (see Cirillo et al., 2019 for review), based on a theory that the right hemisphere is more involved in withdrawal-type emotions such as anxiety (see Zwanzger et al., 2009 for discussion). Overall, hemispheric imbalances have been suggested in several clinical contexts, including MDD (Jiang et al., 2019), bipolar disorder (Moebus, Quirin, and Ehrlenspiel, 2023), and schizophrenia (Oertel-Knöchel and Linden, 2011).

### Conclusion

We discovered and prospectively replicated generalized laterality effects that transcend individual specialized association networks. Although these networks contain bilateral regions that can be co-activated, they can also be functionally dissociated by hemisphere during verbal versus nonverbal processing. Perhaps the time has come to revisit the hemispheric specialization of cognitive functions more deeply, especially since hemisphere-specific targeting is central to therapeutic neuromodulation.

## Materials and Methods

### Overview

Hemispheric lateralization was explored by estimating the organization of networks and then testing the degree to which the networks responded during processing of verbal versus nonverbal materials. In an initial discovery study (*N* = 14), distributed association networks were estimated within each intensively studied individual from resting-state fixation data including identifying all the bilateral components of each network. Task responses were then measured from independent task data within the same participants to explore whether the distinct networks or their lateralized component regions responded differentially. We discovered a hemispheric laterality effect by which processing of verbal versus nonverbal materials differentially modulated the left and right component regions of multiple association networks that span the cortical hierarchy. Given this surprising result, the full study was replicated in an independent study (*N* = 15).

### Participants

For the initial discovery study, paid adult volunteers (*N* = 14; mean age = 22.4, SD = 3.0; 13 right-handed; 8 female) participated in 2-3 MRI sessions. These participants are labeled *S1*-*S14* and a subset of their data (*S1*-*S9*) have been previously reported in a paper focused on the organization of prefrontal cortex (DiNicola, Sun, and Buckner, 2023). For the independent replication study, paid adult volunteers (*N* = 15; mean age = 22.1, SD = 3.9; 15 right-handed; 9 female) participated in 8-11 MRI sessions. These participants are labeled *P1*-*P15* and their data have been reported in papers focused on the cerebral cortex (Du et al., 2024), cerebellum (Saadon-Grosman et al., 2024), striatum (Kosakowski et al., 2024; 2025), hippocampus (Angeli et al., 2025), and thalamus (Du et al., 2025). Here the data are reanalyzed with a novel focus on hemispheric laterality effects.

Participants provided informed consent through a protocol approved by the Harvard University Institutional Review Board.

### MRI Data Acquisition and Processing

Neuroimaging was conducted at the Harvard Center for Brain Science using a 3T Siemens MAGNETOM Prisma^fit^ MRI scanner equipped with a 32-channel head coil (Siemens Healthineers AG, Erlangen, Germany). For the discovery dataset, structural images were acquired with a T1-weighted (T1w) multi-echo magnetization-prepared rapid acquisition gradient-echo (ME-MPRAGE) sequence (van der Kouwe et al., 2008), providing 1.2-mm isotropic voxels (TR = 2,200 ms; TE = 1.57, 3.39, 5.21, 7.03 ms; TI = 1,100 ms; flip angle = 7°; matrix = 192 × 192 × 144; in-plane GRAPPA acceleration factor = 4). For the replication dataset, high-resolution T1w and T2-weighted (T2w) scans were acquired based on the Human Connectome Project sequences (HCP; Harms et al., 2018). T1w ME-MPRAGE parameters: voxel size = 0.8 mm; TR = 2,500 ms; TE = 1.81, 3.60, 5.39, and 7.18 ms; TI = 1,000 ms; flip angle = 8°; matrix = 300 × 320 × 208; in-plane GRAPPA acceleration = 2. T2w sampling perfection with application-optimized contrasts using different flip angle evolution sequence (SPACE) parameters: voxel size = 0.8 mm; TR = 3,200 ms; TE = 564 ms; matrix = 300 × 320 × 208; in-plane GRAPPA acceleration = 2. Rapid T1w structural scans were also obtained as backup using the discovery dataset sequence.

Functional runs for resting-state fixation and all tasks used a multiband gradient-echo echo-planar imaging (EPI) sequence sensitive to the blood oxygenation level-dependent (BOLD) contrast: 2.4-mm isotropic voxels; TR = 1,000 ms; TE = 33 ms; flip angle = 64°; matrix = 92 × 92 × 65. The sequence was generously provided by the Center for Magnetic Resonance Research at the University of Minnesota. During BOLD acquisition, participants’ eye movements were monitored using the EyeLink 1000 Core Plus system with a Long-Range Mount (SR Research, Ottawa, ON, Canada) and assigned a “sleepiness score.” At each session, dual-gradient-echo B0 field maps were acquired: TE = 4.45 and 6.91 ms with slice prescription and spatial resolution matched to the BOLD sequence. All BOLD runs began with 12 sec of fixation for T1 stabilization. Participants viewed stimuli presented on a rear-projected display (F80-4K7 Laser Phosphor Projector; Barco, Kortrijk, Belgium) via a mirror mounted to the head coil. The viewing location was calibrated to ensure a central and comfortable view. Participants were instructed to stay still and alert prior to each scan. Each BOLD fMRI run was examined for quality, and all exclusions were finalized prior to task analysis, as reported previously (DiNicola, Sun, and Buckner, 2023; Du et al., 2024).

Data were processed using the openly available “iProc” pipeline, as previously described (DiNicola, Sun, and Buckner, 2023; Du et al., 2024; see also Braga et al., 2019). iProc maximizes cross-session alignment within individuals and minimizes interpolations to retain as much as possible of the detailed functional anatomy within each individual participant. iProc integrates tools from FreeSurfer (v6.0.0; Fischl, 2012), FSL (v5.0.4; Jenkinson et al., 2012) and AFNI (Cox, 1996; 2012). All data were resampled to the fsaverage6 cortical surface mesh (using trilinear interpolation) and smoothed with a 2-mm full-width-at-half-maximum Gaussian kernel. Analyses were performed on the cortical surface.

### Cortical Network Estimation

Within-individual cortical networks were estimated using a 15-network MS-HBM (Du et al., 2024; see also Kong et al., 2019). Descriptions of key steps are summarized here. Using processed resting-state fixation runs as input, the MS-HBM assigned all cortical surface vertices in fsaverage6 space to one of 15 networks in both hemispheres, identifying all the bilateral regions of each network. For each run, correlations were computed between the time series of each surface vertex and 1,175 uniformly distributed regions across the surface. An initial functional connectivity profile was generated by binarizing the top 10% of these correlations. The MS-HBM was initialized with a 15-network group-level prior derived from the HCP S900 data release and utilized an expectation-maximization algorithm to converge on individual network assignments based on functional connectivity profiles. Participants were run through the model in small groups (see footnote 2 in DiNicola, Sun, and Buckner, 2023), yielding network assignments for all participants in each group. For the discovery study, the same network estimates from DiNicola, Sun, and Buckner (2023) were carried forward for *S1-11* (see “Network Identification” section in DiNicola, Sun, and Buckner, 2023), and additional participants *S12-14* were run through the model as a group. For the replication study, network estimates from Du et al. (2024) for *P1-15* were used.

Nine association networks were examined, spanning multiple levels of the cortical hierarchy and representing a diverse set of functional domains: Default Network A, DN-A; Default Network B, DN-B; Language, LANG; Frontoparietal Network A, FPN-A; Frontoparietal Network B, FPN-B; Cingulo-Opercular, CG-OP; Salience, SAL; Dorsal Attention Network A, dATN-A; Dorsal Attention Network B, dATN-B.

### Left and Right Hemisphere Components of Cortical Networks

For each of the nine networks of interest, left and right hemisphere masks were created. First, the full 15-network estimates were separated into their left and right hemisphere components using Connectome Workbench’s *wb_command -cifti-separate* (v1.3.2; Marcus et al., 2011). The proportion of total vertices (that were assigned a network label) in each hemisphere contained within each network was calculated, as a proxy for the network’s relative size or surface area on the left and the right. In addition, for each individual network, binarized hemisphere-specific masks were created with *wb_command -gifti-label-to-roi* (v1.3.2). These masks were used for quantifying network laterality effects in the task paradigms.

### Task Paradigms and Analysis Within the General Linear Model (GLM)

Participants completed a battery of tasks as described in DiNicola, Sun, and Buckner (2023) and Du et al. (2024). Here the data from the Resting-State Fixation, Sentence Processing, and Working Memory (N-Back) tasks were reanalyzed.

#### Resting-State Fixation Task

Resting-State Fixation data were used for cortical network estimation. During Resting-State Fixation task runs (7 min 2 sec each run), participants fixated on a central black crosshair against an off-white background. Each participant in the discovery dataset completed 5-6 runs, and each participant in the replication dataset completed 17-24 runs.

#### Sentence Processing Task

The Sentence Processing task targeted meaning-based processing within the domain of language. Adapted from Fedorenko and colleagues (2010; 2012), task runs (5 min 0 sec each run) contrasted passively reading strings of words, presented one word at a time, that formed meaningful sentences (Sentence condition) to passively reading strings of nonwords (Nonword condition). Each run included 6 Sentence and 6 Nonword blocks, and each block consisted of 3 strings (12 words/nonwords, 0.45 sec per word), with a cue (0.5 sec) to press a button (right index finger) at the end of each string. Sentence and Nonword blocks were evenly distributed across all order positions, and no more than two consecutive blocks of the same type appeared in a row. Runs also included 4 extended fixation blocks (18 sec). Participants completed 4 runs each in the discovery study and 6 runs each in the replication study (2 participants in replication dataset completed an additional 6 runs of the task). 9 of 14 participants in the discovery study and all participants in the replication study completed the Sentence Processing task.

The analyzed contrast was the Sentence minus Nonword condition. The task GLM included run-level regressors for each condition (Sentence, Nonword), and produced z-statistical maps for Sentence minus Nonword, which were averaged across runs to yield the “Sentence Processing” contrast map. This contrast has been shown to preferentially recruit a left-lateralized association language network (LANG; e.g., see Braga et al., 2020). Models were created using FSL’s first-level Feat (v5.0.4) and used a canonical double-gamma hemodynamic response function and its temporal derivative. Hemisphere-specific network masks were applied to extract the mean z-values for the left and right components of each network.

#### Working Memory (N-Back) Task with Verbal and Nonverbal Stimulus Categories

The “N-Back” task encouraged participants to hold and update items in working memory, including varying the working memory load. The “N-Back Load Effect” (2-Back > 0-Back) has been shown to robustly recruit a frontoparietal network (FPN-A, e.g., see Du et al., 2024). Critical to the focus of this paper, the N-Back task runs (4 min 44 sec each run) included distinct blocks of exclusively verbal and nonverbal stimulus categories (Face, Scene, Letter, Word).

Within N-Back blocks, participants indicated if each stimulus matched either an initial cue (low load 0-Back Condition) or the stimulus seen two before in the sequence (high load 2-Back Condition). Stimulus category differed between blocks. Face stimuli were unfamiliar faces from the HCP (Barch et al., 2013); scene stimuli were provided by the Konkle Laboratory (Josephs and Konkle, 2020; Konkle et al., 2010). Letter stimuli were consonants; word stimuli were 1-syllable words. For the Face, Scene, and Letter categories, a correct match was defined as an identical stimulus (e.g., the same exact face, scene, or letter). For the Word category, a correct match was a rhyming word (e.g., “sound” would be a match with “ground”).

Each run had 8 N-Back task blocks (25 sec) with extended fixation blocks (15 sec each) in-between, occurring after every 2 task blocks. The N-Back block type (0-Back or 2-Back Face, Scene, Letter or Word) was cued with the first stimulus, and each block consisted of 10 stimuli including the cue. Each stimulus was shown for 2 sec, followed by a 0.5-sec intertrial interval. Participants were instructed to respond “Match” (right index finger) or “Non-Match” (left index finger) on each trial following the cue. In each N-Back block, there were 2 targets and 2 non-target lures; positions of targets and lures were counterbalanced within and across runs. In the 2-Back condition, lures were repeated stimuli or rhymes presented in incorrect positions, while in the 0-Back condition, lures were repeated stimuli or rhymes that did not correspond to the cue. The ordering of 0-Back and 2-Back condition and stimulus category were each counterbalanced across runs, and no categories appeared twice in a row. All participants in both the discovery and replication studies completed 8 runs of the N-Back task, except for *P5* who completed 7 runs.

For analysis of the N-Back task, each block of trials (including the cue) was modeled as a separate regressor. Models were again created using FSL’s first-level Feat (v5.0.4) and used a canonical double-gamma hemodynamic response function and its temporal derivative. Each load x category pairing was computed as its own map for every participant (2-Back Face, 0-Back Face, 2-Back Word, 0-Back Word, etc). Hemisphere-specific network masks were applied to extract the mean z-values for the left and right components of each network for each of the critical contrasts. To estimate the networks active during the N-Back Load Effect generally, the 2-Back > 0-Back contrast was modeled combining stimulus conditions. To specifically examine the effects of stimulus category, the N-Back Face > Word contrast was estimated as described in the next section.

### Explorations of Verbal versus Nonverbal Processing

To examine the extent to which verbal versus nonverbal materials are processed differentially by cortical networks, the critical analysis was the Face > Word task contrast in the N-Back task. This contrast was calculated within each individual by holding load level constant, computing the 2-Back Face minus 2-Back Word and 0-Back Face minus 0-Back Word maps, and then averaging them to yield a single best estimate of the effect of the stimulus category. To quantify the hemispheric effects, mean z-values for each network were extracted separately for the left and right hemispheres using the corresponding hemispheric masks. This yielded 9 (networks) by 2 (hemisphere) values per participant. The critical analyses focused on whether networks showed a hemispheric laterality effect for the N-Back Face > Word task contrast. A repeated measures ANOVA with hemisphere as the within-subject factor, and mean z-value as the dependent variable was conducted. Post-hoc analyses, reported in the Supplemental Materials, also explored effects that include the Letter and Scene stimulus categories.

### Prospective Replication Analysis of Generalized Laterality Effects

After analyzing the N-Back Face > Word task contrast from the discovery study, we observed hemispheric laterality effects that spanned multiple networks (see **Figure 3**). Given this was an unexpected result, we sought to prospectively replicate this effect in an independent data set. The replication study data were set aside and not examined until the present approach was defined and tested for replication.

We selected four networks to test for replication. The four networks were selected strategically to establish the generality of the effect. Within higher-order association cortex there are distinct networks linked to domain-specialized processing and domain-flexible processing associated with cognitive control (Du et al., 2024; see also Nee, 2021). From this group of networks, we selected LANG, which is well established to be domain-specialized for processing within the domain of language (Fedorenko, Ivanova, and Regev, 2024), and FPN-A, which responds robustly to effortful task demands flexibly across a range of domains (Duncan, 2010; Badre and Nee, 2018). Both networks showed significant laterality effects in the discovery study. From among candidate second-order association networks, we selected CG-OP, also known as the Action-Mode Network, which participates in motor preparation (Dosenbach, Raichle, and Gordon, 2025), and dATN-A, a network that participates in external attention (Corbetta and Shulman, 2002). Both networks also showed significant laterality effects in the discovery study.

Anatomically, LANG and FPN-A populate the distributed association zones across the cortex including prefrontal, posterior parietal, and temporal association cortex. CG-OP has prominent components juxtaposed with somatosensory and motor cortices; dATN-A is anatomically juxtaposed with retinotopic visual cortex (see flatmap representations in Figs. 17-19 of Du et al., 2024). Thus, these networks display a range of topographic patterns that span the association cortices.

By targeting four replication networks that participate in diverse functions and that are anatomically distinct we sought to rigorously test the hypothesis that the hemispheric laterality effect generalizes across multiple association networks. This is a stringent replication test: to confirm the effect requires four separate tests to each be significant. Specifically, we hypothesized there would be significantly greater activity in the right than left hemisphere for the N-Back Face > Word task contrast using paired t-tests in all four targeted networks (LANG, FPN-A, CG-OP, and dATN-A). To support the hypothesis that the hemispheric laterality effect generalizes, the four tests each are required to be significant (*p* < 0.05 one-tailed). Thus, the chance probability is considerably lower than the individual test threshold^2^.

## Acknowledgements

We thank the Harvard Center for Brain Science neuroimaging core and FAS Division of Research Computing. We thank Tim O’Keefe and Daniel Asay for assisting in optimization of data processing, Ross Mair for MRI physics support, Theo Tobel for assistance with proofreading, Jingnan Du, Anne Billot, Mark Eldaief, Vaibhav Tripathi, Xiangyu Wei, Max Elliott, Heather Kosakowski, and Peter Angeli for helpful feedback and discussion.

This work was supported by NIH grant MH124004 (R.L.B.), NIH grant T32AG000222 (W.S.), NIH Shared Instrumentation grant S10OD020039 (R.L.B.), Kent and Liz Dauten (R.L.B.), and the Paul and Daisy Soros Foundation (W.S.). During data acquisition, L.M.D. was supported by NSF Graduate Research Fellowship Program Grant DGE1745303.

## Declaration of Interests

The authors declare no competing interests.

## Footnotes

1 Frontoparietal Network B (FPN-B) is the most strongly right-lateralized association network (Braga et al., 2020). FPN-B’s response properties have been an enigma with an attenuated response relative to FPN-A (Du et al., 2024). When we began this line of work, our hypothesis was that FPN-B might be specialized for nonverbal processing. As the results will reveal, our initial hypothesis was not supported. Given the surprising results we undertook a full prospective replication of the novel hypotheses that emerged. Note also that the network labeled FPN-B here, which is labeled using the naming convention of Xue et al. (2021), is referred to as FPN-A in Braga et al. (2020).

2 The hypothesis of generality requires that *each* of the four tests be significant (leading to chance probability being much more conservative than the individual *p* < 0.05 threshold used for each separate test). In practice, the results were sufficiently robust that they would also withstand multiple comparison correction, even for the full set of all four networks (all were *p* < 0.001). However, multiple comparison correction is not formally required in a dependent set of hypotheses as targeted here so we outline the rationale and use the *a priori* test and statistical threshold as established in advance of the results.

## Notes

### Competing Interest Statement

The authors have declared no competing interest.

### Summary of Updates

Minor copy-editing changes; Methods section updated to clarify network vertex calculations and participant details; updated references; supplemental figures now included.

