## Supplemental Figures for "Verbal versus Nonverbal Processing Leads to Generalized Hemispheric Laterality Effects that Span Multiple Networks"

This PDF file includes:

**Supplemental Figures 1 to 13**

**The Supplemental Figures describe results including: (1) network estimates in one representative participant from the replication study, (2-3) plots showing the mean percentage of vertices in each association network for both left and right hemispheres in the discovery and replication studies, (4-5) plots showing the percentage of vertices in each association network for both left and right hemispheres in every individual in the discovery and replication studies, (6-7) plots showing task-specific effects in association networks in the discovery and replication studies, (8-10) plots of the N-Back Face > Letter, Scene > Word, and Face > Word task contrasts in the discovery dataset, (11-13) plots of the N-Back Face > Letter, Scene > Word, and Face > Word task contrasts in the replication dataset.**

**Supplemental Figure 1. Full cortical network estimates in one representative individual (replication dataset).** Mirroring Figure 1, network estimates are illustrated for a representative individual from the replication dataset. Network name abbreviations: DN-A, Default Network A; DN-B, Default Network B; LANG, Language; FPN-A, Frontoparietal Network A; FPN-B, Frontoparietal Network B; CG-OP, Cingulo-Opercular; SAL, Salience; dATN-A, Dorsal Attention A; dATN-B, Dorsal Attention B.

**Supplemental Figure 2. Mean percentage of vertices in each association network for both left and right hemispheres (discovery dataset).** This plot displays the % of total labeled vertices in the left and right hemispheres included in each of 9 association networks as a proxy for the relative size / surface area occupied by each cortical network. Displayed is the mean across the 14 participants from the discovery dataset. Error bars show standard error of the mean. Direct comparison of the number of vertices between left and right hemispheres indicates that the LANG network is left-lateralized on average in the group. By contrast, FPN-B is right-lateralized on average in the group. However, despite varying degrees of lateralization, each network still has regions in both hemispheres.

**Supplemental Figure 3. Mean percentage of vertices in each association network for both left and right hemispheres (replication dataset).** Following the same steps as Figure S2, this plot shows the % of total labeled vertices in the left and right hemispheres included in each network. Displayed is the mean across the 15 participants from the replication dataset. Error bars show standard error of the mean. Again, LANG is left-lateralized, while FPN-B is right-lateralized. Despite varying degrees of lateralization, each network has bilateral regions.

**Supplemental Figure 4. The percentage of network vertices in the left and right hemispheres plotted for each individual in the discovery dataset**. Plots display the % of labeled vertices in each hemisphere contained within the DN-A, DN-B, LANG, FPN-A, FPN-B, CG-OP, SAL, and dATN-A networks in the discovery dataset. Lighter-shaded bars represent the left hemisphere, and darker-shaded bars represent the right hemisphere. Critically, each participant has bilateral regions in each network.

**Supplemental Figure 5. The percentage of network vertices in the left and right hemispheres plotted for each individual in the replication dataset.** Plots display the % of labeled vertices in each hemisphere contained within the DN-A, DN-B, LANG, FPN-A, FPN-B, CG-OP, SAL, and dATN-A networks in the replication dataset. Lighter-shaded bars represent the left hemisphere, and darker-shaded bars represent the right hemisphere. As shown in the discovery dataset, each participant has bilateral regions in each network.

**Supplemental Figure 6. Association networks respond in a task-specific manner (discovery dataset).** Each plot shows the mean response for a given task contrast in the discovery study, calculated as mean z-values within the boundaries of association networks, with the hemispheres combined. Error bars show standard error of the mean. Dotted lines denote the four groups (**I-IV**) of networks, depicted in Figures 1 and S1. (**Top**) The Sentence Processing task contrast preferentially recruits the LANG network. (**Bottom**) Although broader than the language localizer, the N-Back Load Effect task contrast strongly drives activity in FPN-A and dATN-A, and to a lesser extent FPN-B and SAL. Together, these data show that activity within association networks can be preferentially and differentially modulated by tasks. Asterisks above bars indicate significant positive responses. *** = $p < 0.001$.

**Supplemental Figure 7. Association networks respond in a task-specific manner (replication dataset).** Each plot shows the mean response for a given task contrast in the replication study, calculated as mean z-values within the boundaries of association networks, with the hemispheres combined. Error bars show standard error of the mean. (**Top**) Replication of preferential recruitment of LANG by the Sentence Processing task contrast. (**Bottom**) Replication of robust FPN-A and dATN-A activity in the N-Back Load Effect task contrast and lesser effect in FPN-B and SAL. These data reinforce the finding that different tasks can preferentially modulate activity within cortical networks. Asterisks above bars indicate significant positive responses. *** = $p < 0.001$.

**Supplemental Figure 8. N-Back Face > Letter task contrast functionally dissociates the left and right hemispheres of multiple networks (discovery dataset).** (**Top**) Mirroring the methods used for Figure 4, the discovery dataset's group averaged Face (nonverbal; red/yellow) > Letter (verbal; blue) task contrast is visualized as z-statistical maps on the inflated cortical surface. L, letters; F, faces. (**Bottom**) Mirroring the methods used for Figure 3, this plot shows the mean response for the Face (nonverbal) > Letter (verbal) task contrast, calculated as mean z-values within each hemisphere of association networks. Lighter-shaded bars represent the left (L) hemisphere, and darker-shaded bars represent the right (R) hemisphere. Error bars show standard error of the mean. Bracketed asterisks indicate that the direct comparisons between the left and right hemisphere responses are significantly different. Asterisks by the bars indicate the individual bar's response is significantly different from zero (positive in the right hemisphere, negative in the left hemisphere). * = $p < 0.05$, ** = $p < 0.01$, *** = $p < 0.001$.

**Supplemental Figure 9**. **N-Back Scene > Word task contrast functionally dissociates the left and right hemispheres of multiple networks (discovery dataset).** (**Top**) Mirroring the methods used for Figure 4, the discovery dataset's group averaged Scene (nonverbal; red/yellow) > Word (verbal; blue) task contrast is visualized as z-statistical maps on the inflated cortical surface. W, words; S, scenes. (**Bottom**) Mirroring the methods used for Figure 3, this plot shows the mean response for the Scene (nonverbal) > Word (verbal) task contrast, calculated as mean z-values within each hemisphere of association networks. Lighter-shaded bars represent the left (L) hemisphere, and darker-shaded bars represent the right (R) hemisphere. Error bars show standard error of the mean. Bracketed asterisks indicate that the direct comparisons between the left and right hemisphere responses are significantly different. Asterisks by the bars indicate the individual bar's response is significantly different from zero (positive in the right hemisphere, negative in the left hemisphere). * = $p < 0.05$, ** = $p < 0.01$, *** = $p < 0.001$.

**Supplemental Figure 10. N-Back Face > Word task contrast functionally dissociates the left and right hemispheres of multiple networks (discovery dataset).** Data from Figures 3 and 4 are replotted here to facilitate comparison with the task contrasts in Figures S8 and S9. (**Top**) The discovery dataset's group averaged Face (nonverbal; red/yellow) > Word (verbal; blue) task contrast is visualized as z-statistical maps on the inflated cortical surface. W, words; F, faces. (**Bottom**) This plot shows the mean response for the Face (nonverbal) > Word (verbal) task contrast, calculated as mean z-values within each hemisphere of association networks. * = $p < 0.05$, ** = $p < 0.01$, *** = $p < 0.001$.

**Supplemental Figure 11. N-Back Face > Letter task contrast functionally dissociates the left and right hemispheres of multiple networks (replication dataset).** (**Top**) Mirroring the methods used for Figure 7, the replication dataset's group averaged Face (nonverbal; red/yellow) > Letter (verbal; blue) task contrast is visualized as z-statistical maps on the inflated cortical surface. L, letters; F, faces. (**Bottom**) Mirroring the methods used for Figure 6, this plot shows the mean response for the Face (nonverbal) > Letter (verbal) task contrast, calculated as mean z-values within each hemisphere of four selected cortical networks in groups **I-IV**: LANG, FPN-A, CG-OP, and dATN-A. Lighter-shaded bars represent the left hemisphere, and darker-shaded bars represent the right hemisphere. Error bars show standard error of the mean. Bracketed asterisks indicate that the direct comparisons between the left and right hemisphere responses are significantly different. Asterisks by the bars indicate the individual bar's response is significantly different from zero (positive in the right hemisphere, negative in the left hemisphere). ** = $p < 0.01$, *** = $p < 0.001$.

**Supplemental Figure 12. N-Back Scene > Word task contrast functionally dissociates the left and right hemispheres of multiple networks (replication dataset).** (**Top**) Mirroring the methods used for Figure 7, the replication dataset's group averaged Scene (nonverbal; red/yellow) > Word (verbal; blue) task contrast is visualized as z-statistical maps on the inflated cortical surface. W, words; S, scenes. (**Bottom**) Mirroring the methods used for Figure 6, this plot shows the mean response for the Scene (nonverbal) > Word (verbal) task contrast, calculated as mean z-values within each hemisphere of four selected cortical networks in groups **I-IV**: LANG, FPN-A, CG-OP, and dATN-A. Lighter-shaded bars represent the left hemisphere, and darker-shaded bars represent the right hemisphere. Error bars show standard error of the mean. Bracketed asterisks indicate that the direct comparisons between the left and right hemisphere responses are significantly different. Asterisks by the bars indicate the individual bar's response is significantly different from zero (positive in the right hemisphere, negative in the left hemisphere). ** = $p < 0.01$, *** = $p < 0.001$.

**Supplemental Figure 13. N-Back Face > Word task contrast functionally dissociates the left and right hemispheres of multiple networks (replication dataset).** Data from Figures 6 and 7 are replotted here to facilitate comparison with the task contrasts in Figures S11 and S12. (**Top**) The replication dataset's group averaged Face (nonverbal; red/yellow) > Word (verbal; blue) task contrast is visualized as z-statistical maps on the inflated cortical surface. W, words; F, faces. (**Bottom**) This plot shows the mean response for the Face (nonverbal) > Word (verbal) task contrast, calculated as mean z-values within each hemisphere of *a priori* selected networks. * = $p < 0.05$, ** = $p < 0.01$, *** = $p < 0.001$.

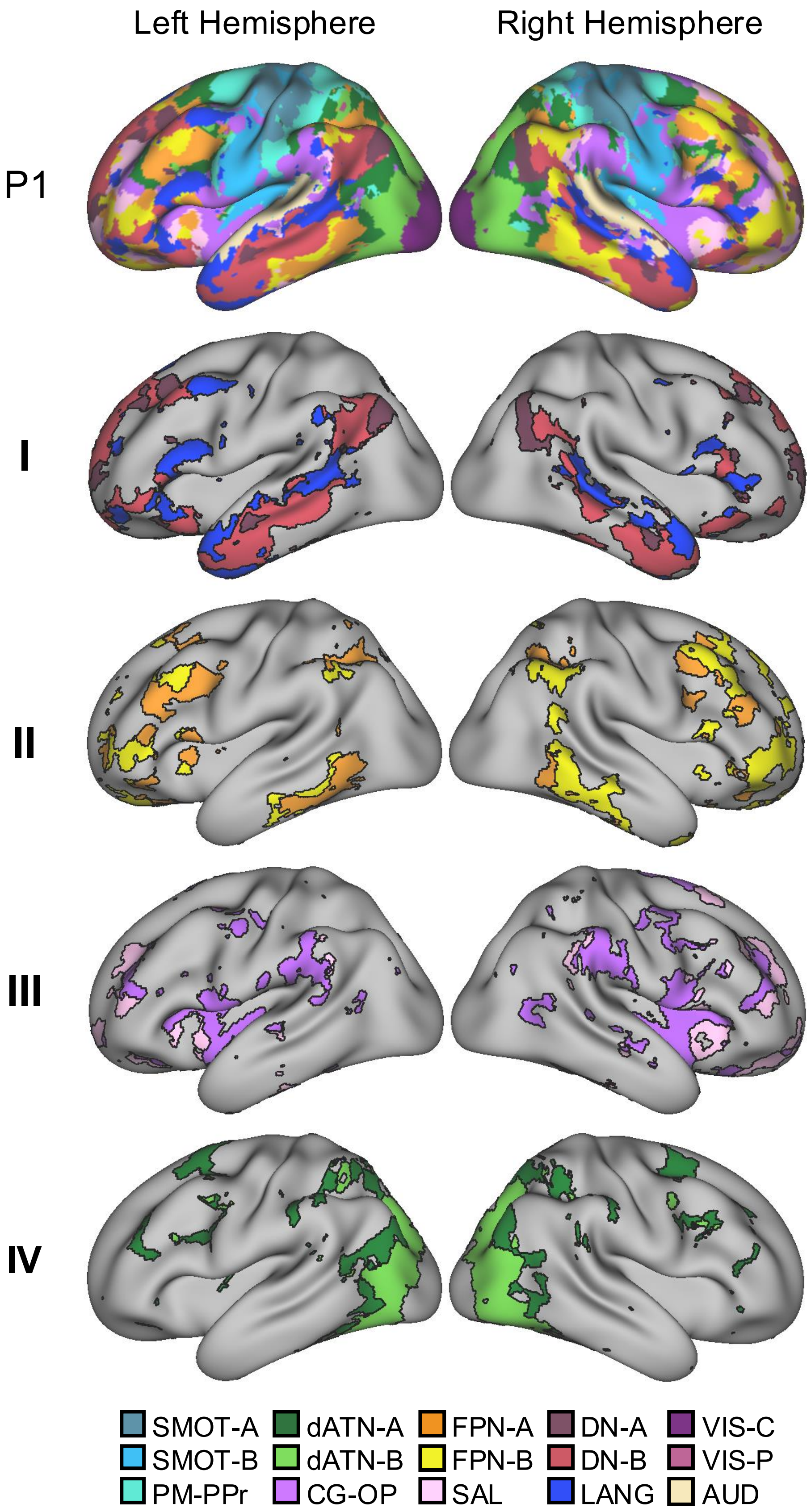

Left Hemisphere
Right Hemisphere
P1
I
II
III
IV
SMOT-A
SMOT-B
PM-PPr
dATN-A
dATN-B
CG-OP
FPN-A
FPN-B
SAL
DN-A
DN-B
LANG
VIS-C
VIS-P
AUD


Figure S1

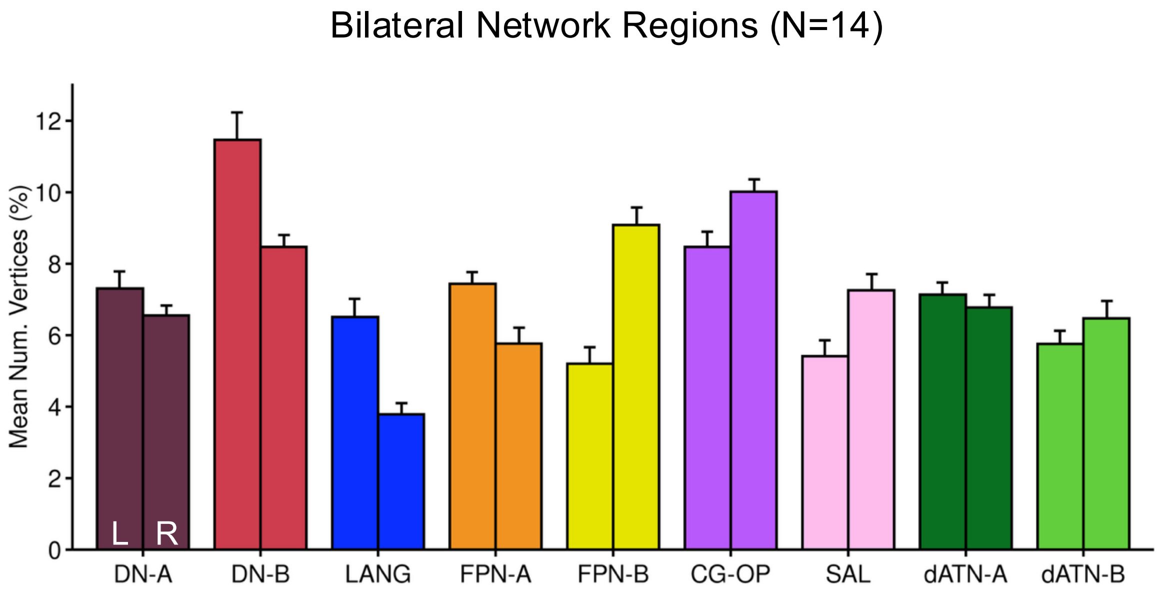

Bilateral Network Regions (N=14)
Mean Num. Vertices (%)
0
2
4
6
8
10
12
L
R
DN-A
DN-B
LANG
FPN-A
FPN-B
CG-OP
SAL
dATN-A
dATN-B


Figure S2

Bilateral Network Regions (N=15)

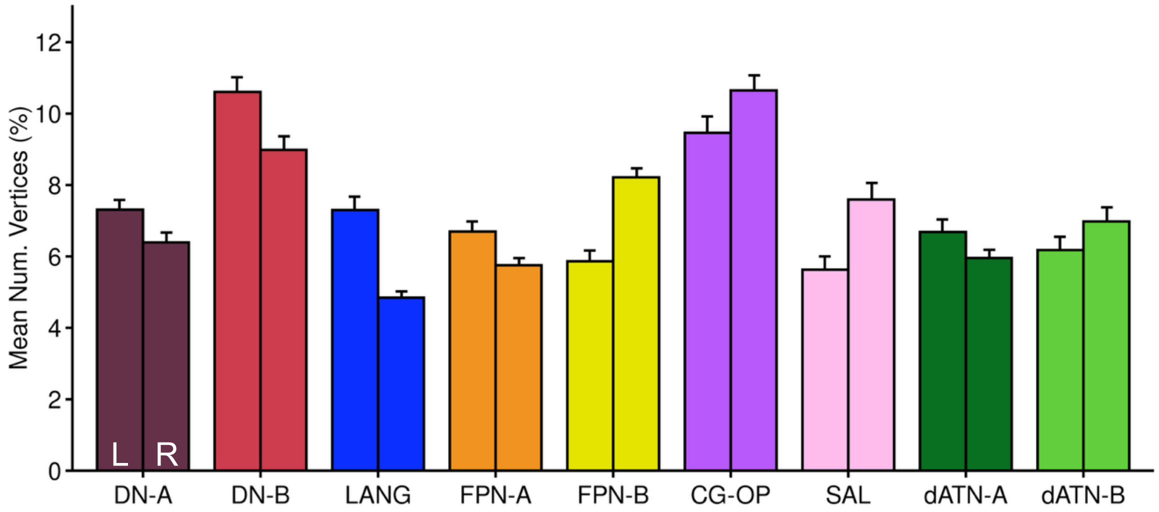


Figure S3

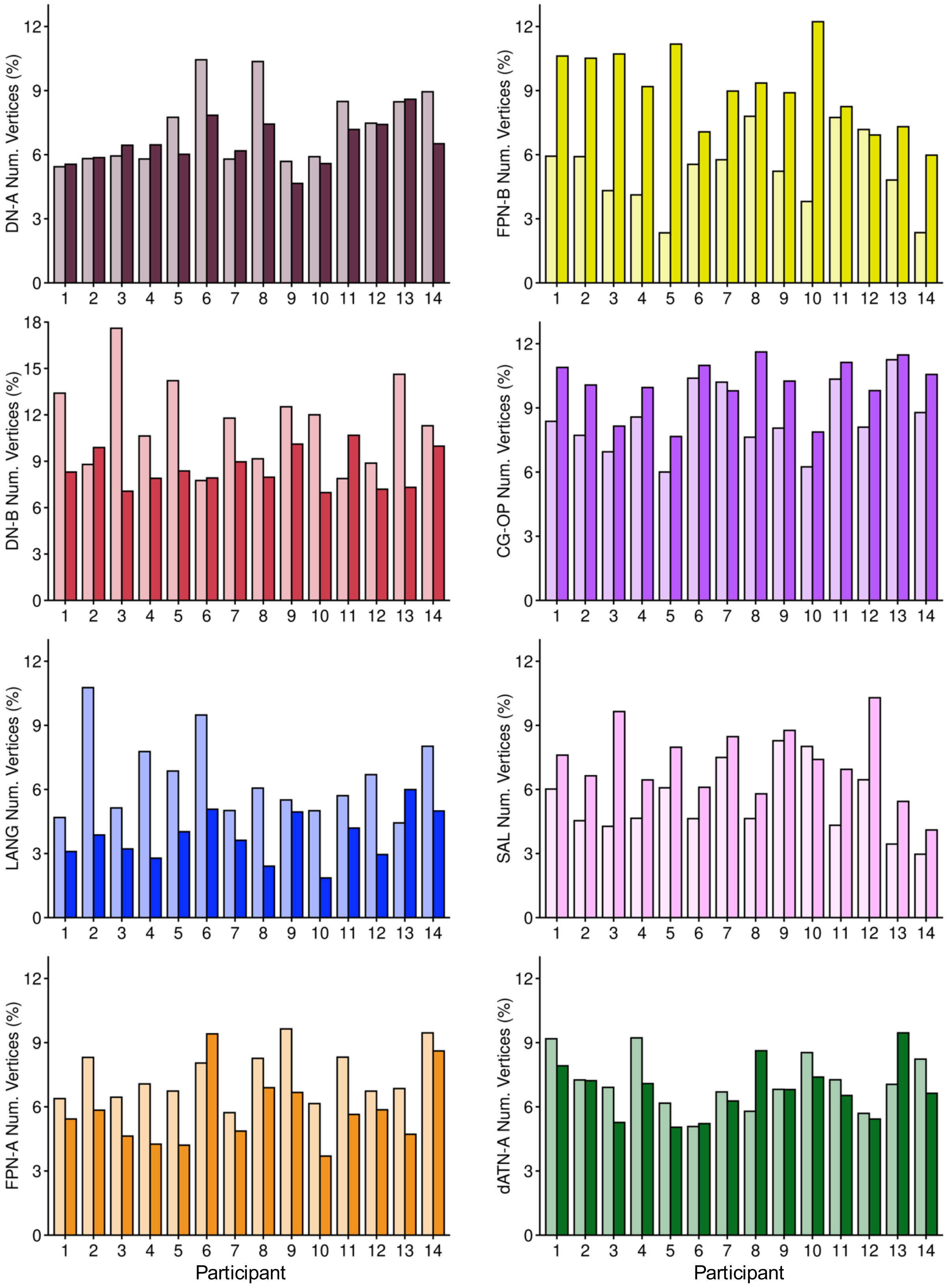
DN-A Num. Vertices (%)
FPN-B Num. Vertices (%)
DN-B Num. Vertices (%)
CG-OP Num. Vertices (%)
LANG Num. Vertices (%)
SAL Num. Vertices (%)
FPN-A Num. Vertices (%)
dATN-A Num. Vertices (%)
Participant
Participant

Figure S4

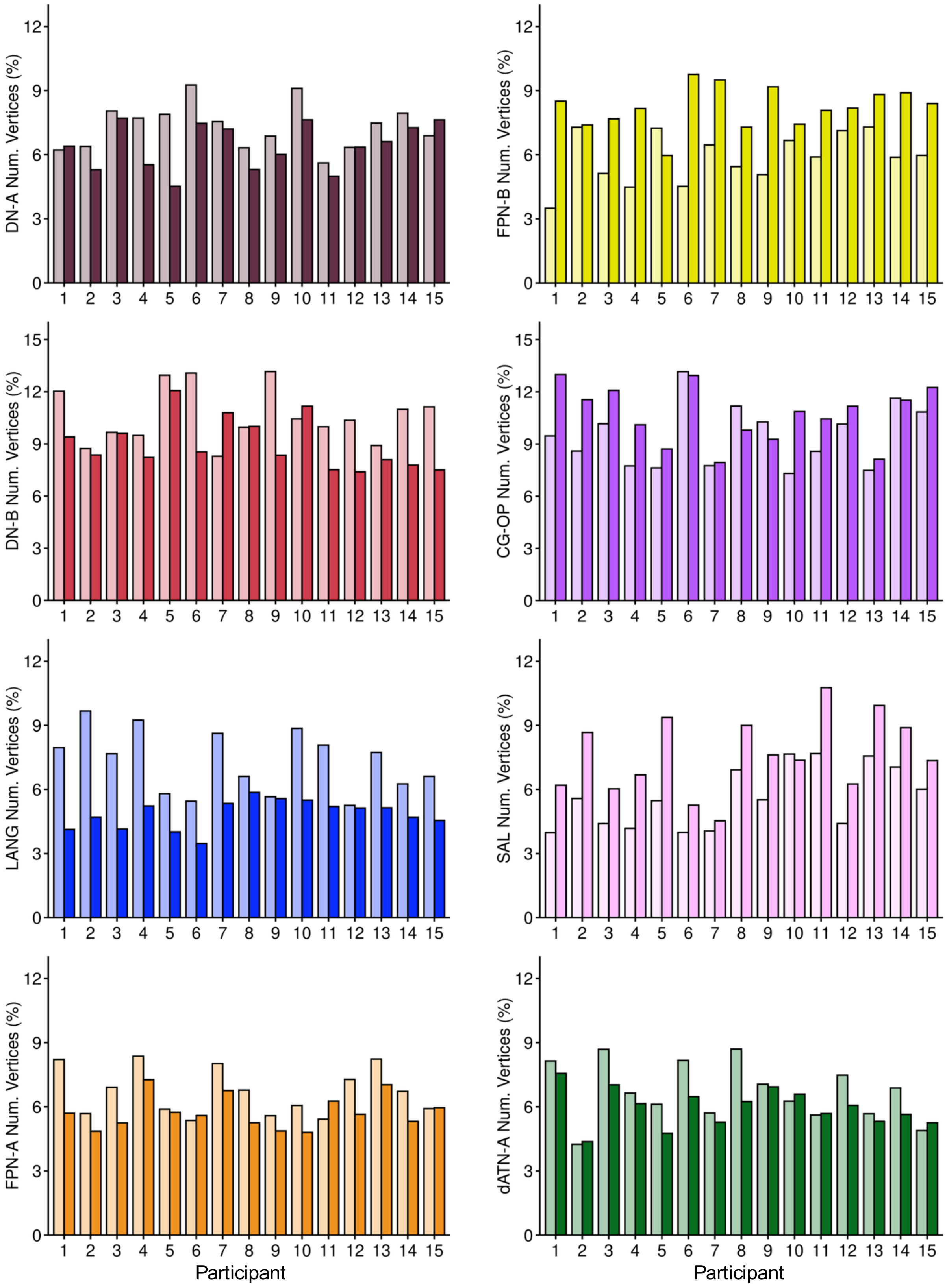
DN-A Num. Vertices (%)
FPN-B Num. Vertices (%)
DN-B Num. Vertices (%)
CG-OP Num. Vertices (%)
LANG Num. Vertices (%)
SAL Num. Vertices (%)
FPN-A Num. Vertices (%)
dATN-A Num. Vertices (%)
Participant
Participant

Figure S5

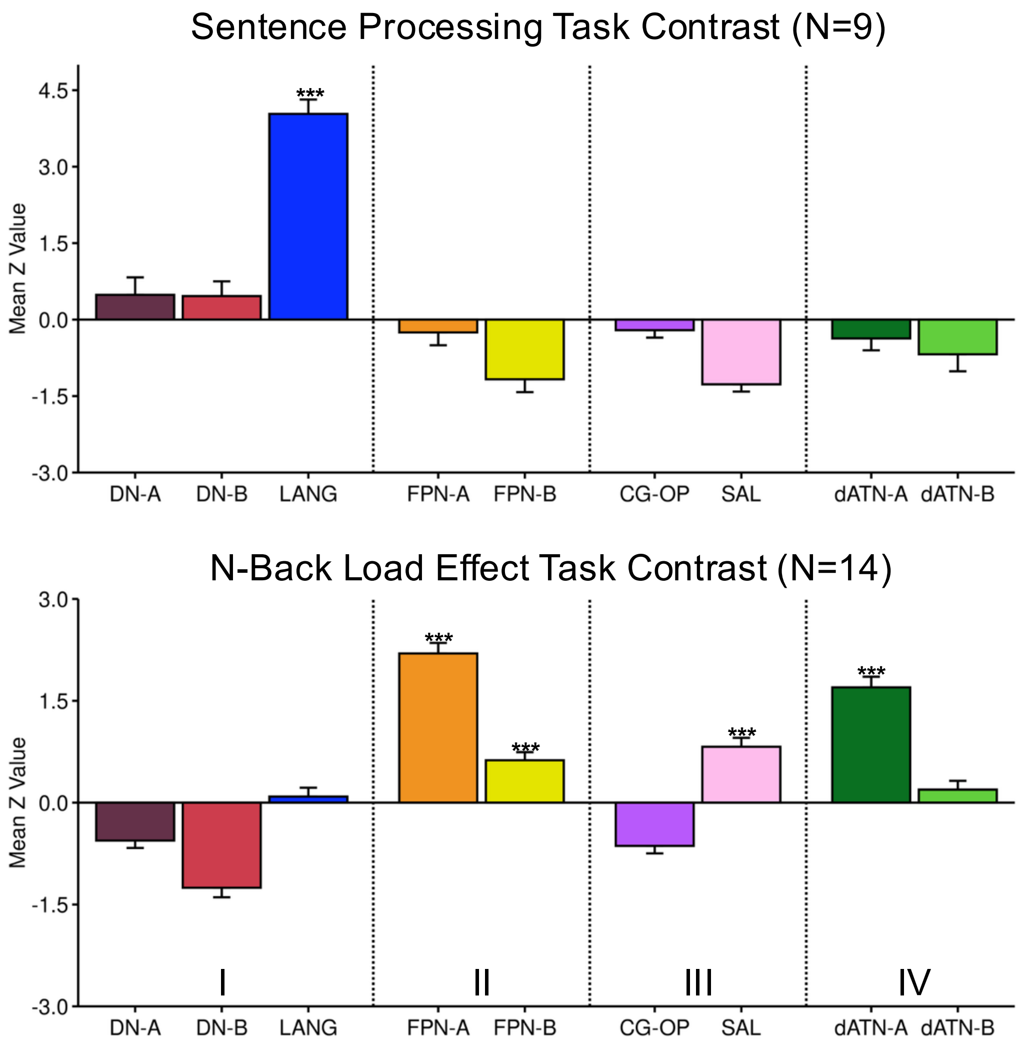

Sentence Processing Task Contrast (N=9)
Mean Z Value
4.5
3.0
1.5
0.0
-1.5
-3.0
***
DN-A
DN-B
LANG
FPN-A
FPN-B
CG-OP
SAL
dATN-A
dATN-B
N-Back Load Effect Task Contrast (N=14)
Mean Z Value
3.0
1.5
0.0
-1.5
-3.0
***
***
***
***
I
II
III
IV
DN-A
DN-B
LANG
FPN-A
FPN-B
CG-OP
SAL
dATN-A
dATN-B


Figure S6

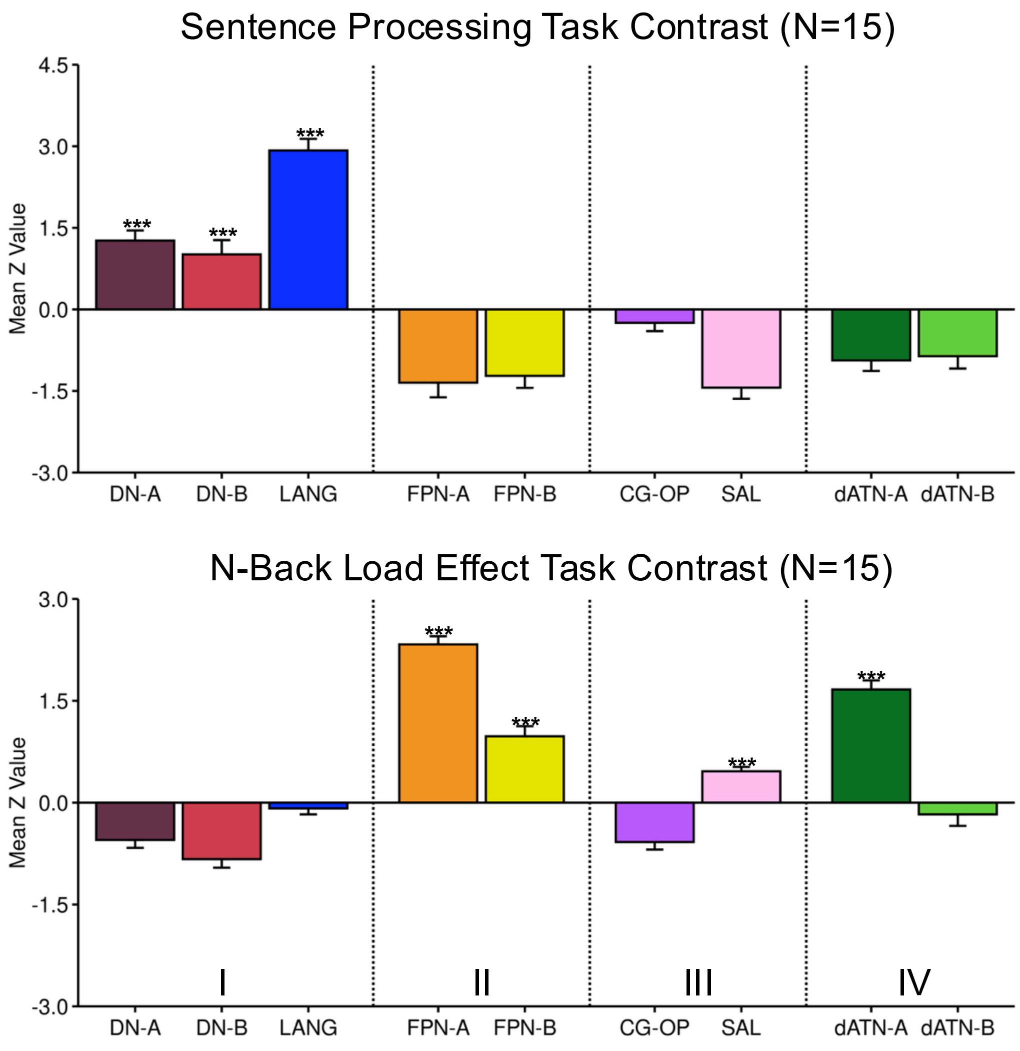

Sentence Processing Task Contrast (N=15)
4.5
3.0
1.5
0.0
-1.5
-3.0
Mean Z Value
***
***
***
DN-A
DN-B
LANG
FPN-A
FPN-B
CG-OP
SAL
dATN-A
dATN-B
N-Back Load Effect Task Contrast (N=15)
3.0
1.5
0.0
-1.5
-3.0
Mean Z Value
***
***
***
***
I
II
III
IV
DN-A
DN-B
LANG
FPN-A
FPN-B
CG-OP
SAL
dATN-A
dATN-B


Figure S7

N-Back Face > Letter Task Contrast
Discovery Group Average

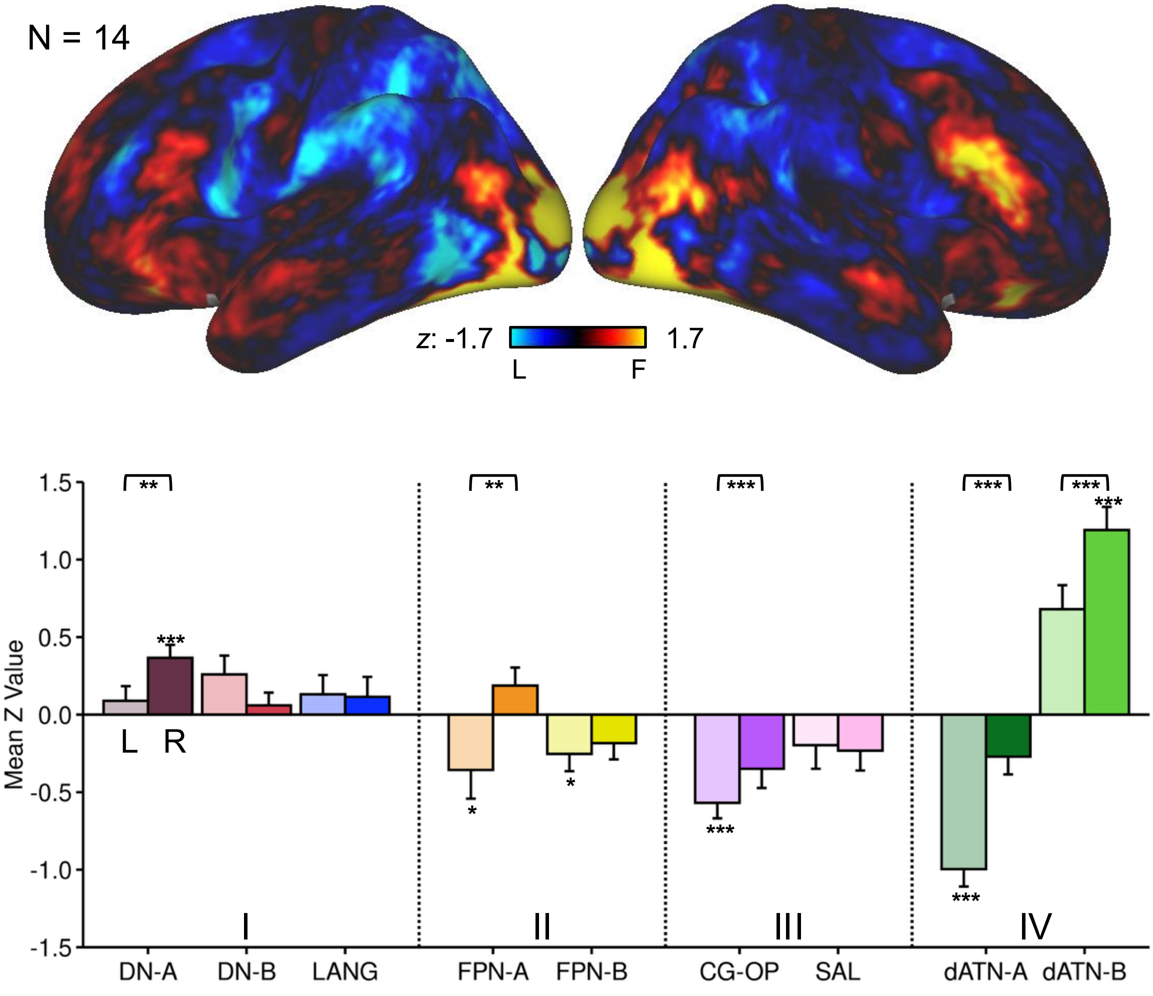


Figure S8

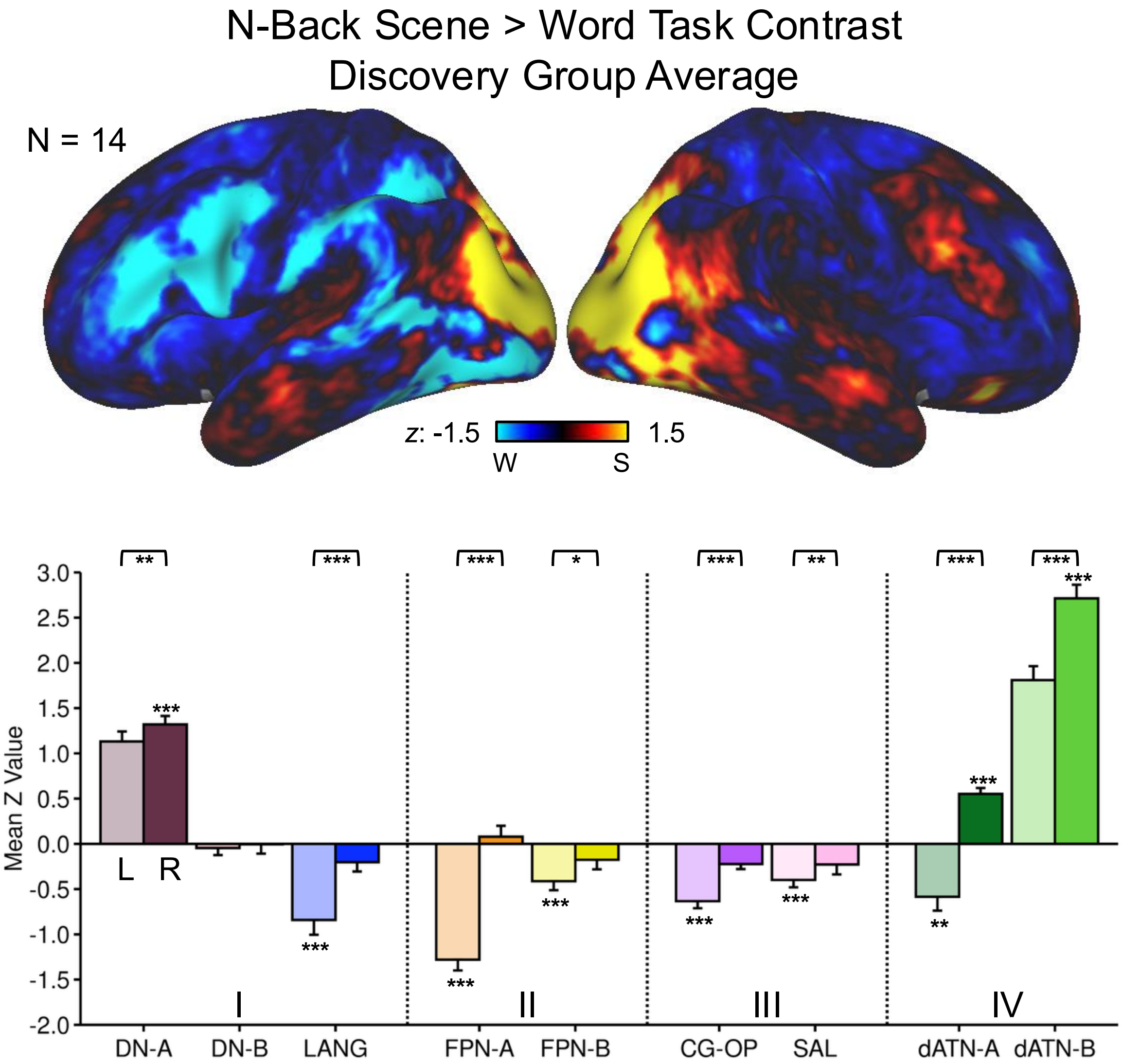

N-Back Scene > Word Task Contrast
Discovery Group Average
N = 14
z: -1.5
1.5
W
S
Mean Z Value
3.0
2.5
2.0
1.5
1.0
0.5
0.0
-0.5
-1.0
-1.5
-2.0
L
R
I
II
III
IV
DN-A
DN-B
LANG
FPN-A
FPN-B
CG-OP
SAL
dATN-A
dATN-B


Figure S9

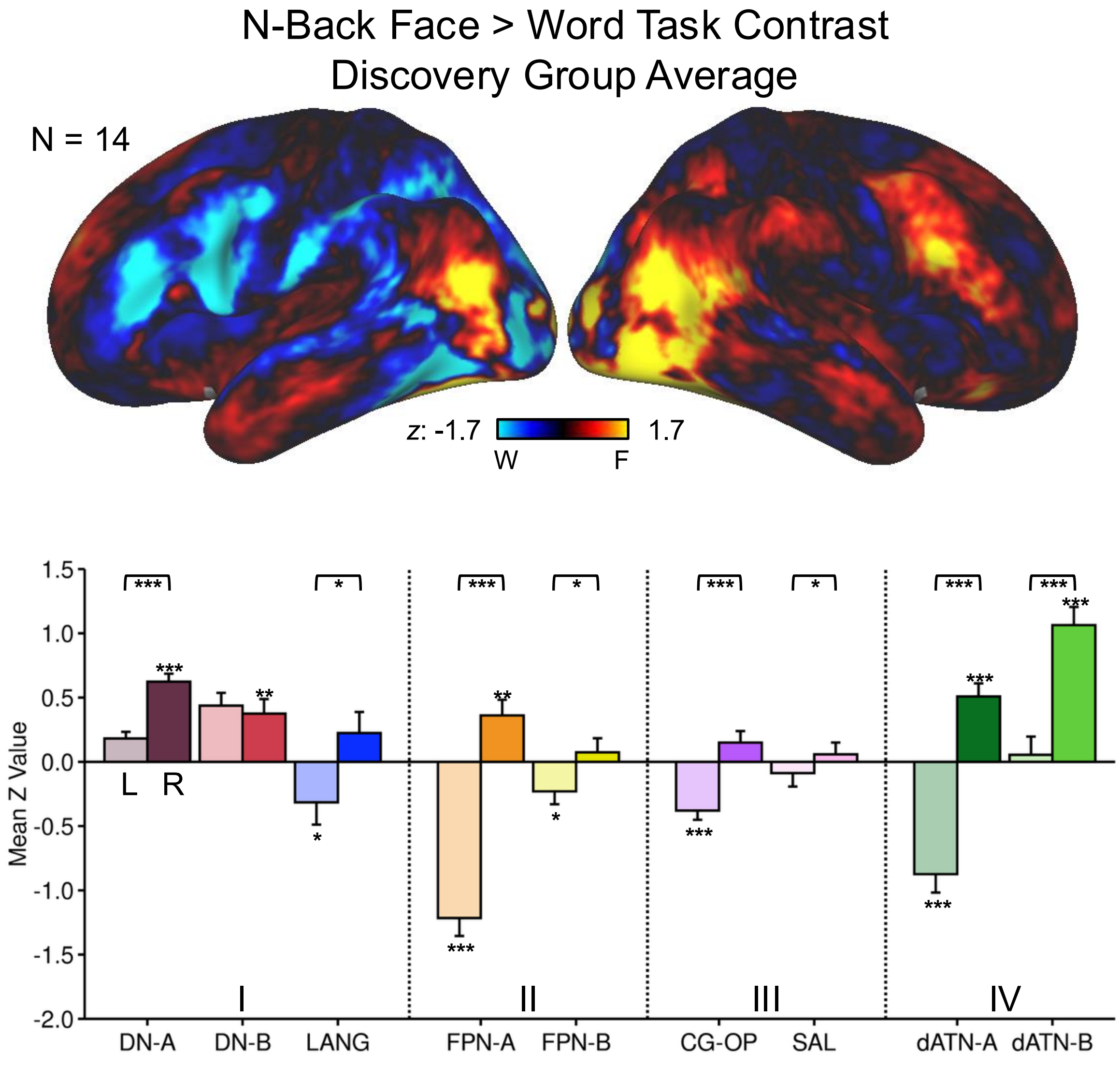
N-Back Face > Word Task Contrast
Discovery Group Average
N = 14
z: -1.7
1.7
W
F
Mean Z Value
1.5
1.0
0.5
0.0
-0.5
-1.0
-1.5
-2.0
L
R
I
II
III
IV
DN-A
DN-B
LANG
FPN-A
FPN-B
CG-OP
SAL
dATN-A
dATN-B

Figure S10

N-Back Face > Letter Task Contrast
Replication Group Average

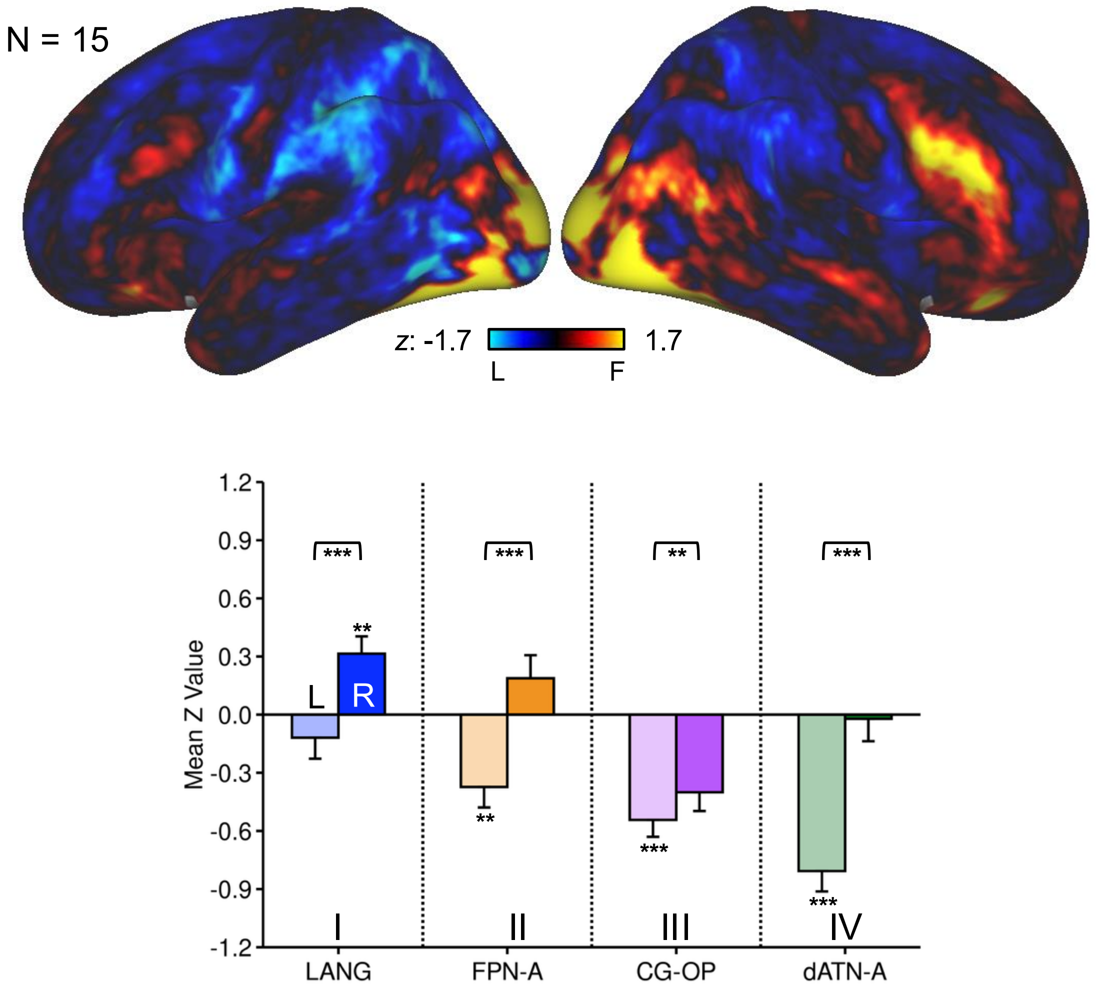


Figure S11

N-Back Scene > Word Task Contrast
Replication Group Average

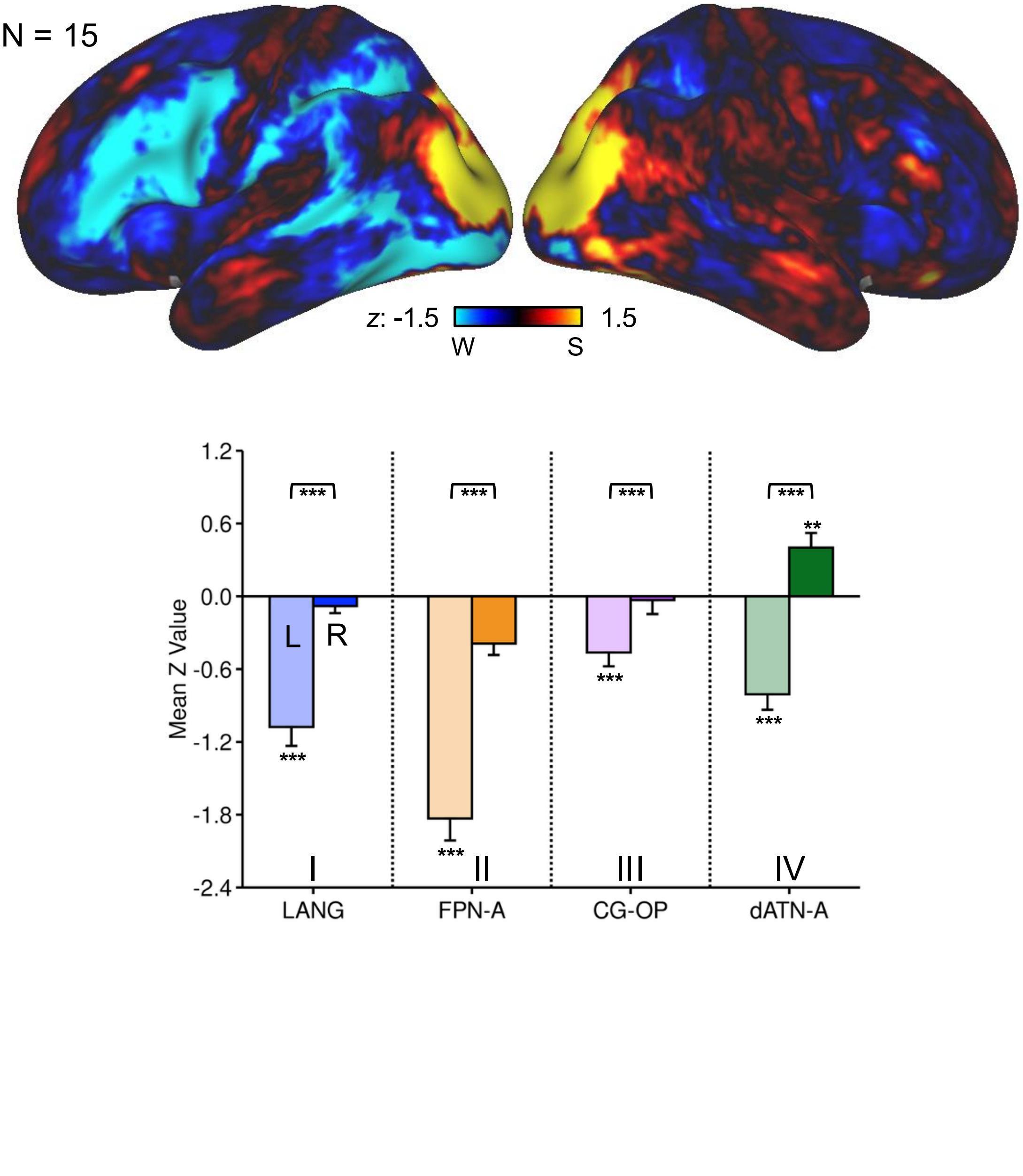


Figure S12

N-Back Face > Word Task Contrast
Replication Group Average

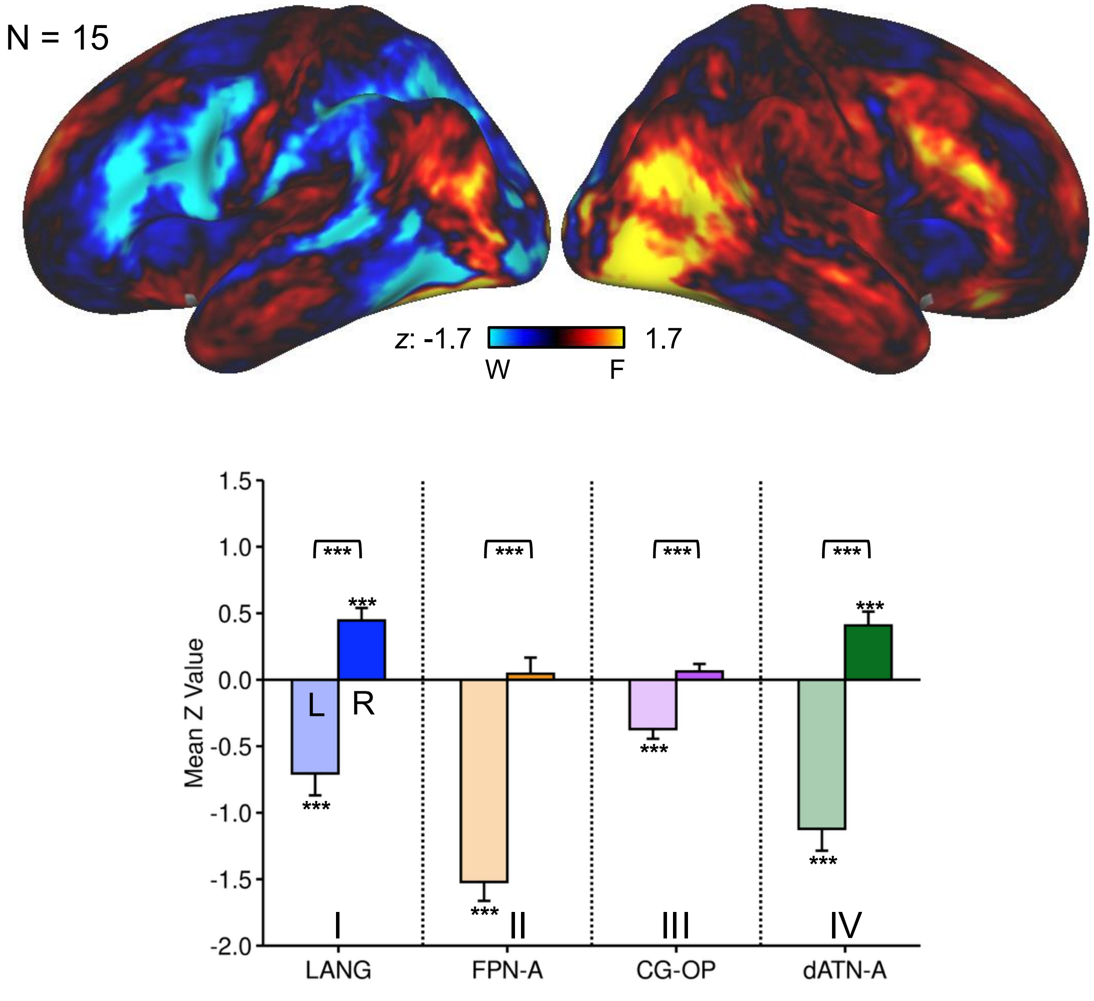


Figure S13
